# Cross-reactive sarbecovirus antibodies induced by mosaic RBD-nanoparticles

**DOI:** 10.1101/2025.01.02.631145

**Authors:** Chengcheng Fan, Jennifer R. Keeffe, Kathryn E. Malecek, Alexander A. Cohen, Anthony P. West, Viren A. Baharani, Annie V. Rorick, Han Gao, Priyanthi N.P. Gnanapragasam, Semi Rho, Jaasiel Alvarez, Luisa N. Segovia, Theodora Hatziioannou, Paul D. Bieniasz, Pamela J. Bjorkman

## Abstract

Broad immune responses are needed to mitigate viral evolution and escape. To induce antibodies against conserved receptor-binding domain (RBD) regions of SARS-like betacoronavirus (sarbecovirus) spike proteins that recognize SARS-CoV-2 variants of concern and zoonotic sarbecoviruses, we developed mosaic-8b RBD-nanoparticles presenting eight sarbecovirus RBDs arranged randomly on a 60-mer nanoparticle. Mosaic-8b immunizations protected animals from challenges from viruses whose RBDs were matched or mismatched to those on nanoparticles. Here, we describe neutralizing mAbs isolated from mosaic-8b–immunized rabbits, some on par with Pemgarda, the only currently FDA-approved therapeutic mAb. Deep mutational scanning, *in vitro* selection of spike resistance mutations, and single-particle cryo-electron microscopy structures of spike-antibody complexes demonstrated targeting of conserved RBD epitopes. Rabbit mAbs included critical D-gene segment RBD-recognizing features in common with human anti-RBD mAbs, despite rabbit genomes lacking an equivalent human D-gene segment, thus demonstrating that the immune systems of humans and other mammals can utilize different antibody gene segments to arrive at similar modes of antigen recognition. These results suggest that animal models can be used to elicit anti-RBD mAbs with similar properties to those raised in humans, which can then be humanized for therapeutic use, and that mosaic RBD-nanoparticle immunization coupled with multiplexed screening represents an efficient way to generate and select broadly cross-reactive therapeutic pan-sarbecovirus and pan-SARS-CoV-2 variant mAbs.

**Significance Statement:** SARS-CoV-2 variants and potential zoonotic sarbecovirus infections continue to threaten human health. Anti-SARS-CoV-2 mAbs that recognize conserved epitopes could be used prophylactically or therapeutically. We present approaches to elicit and identify cross-reactive mAbs using immunizations in animals with mosaic RBD-nanoparticles. We show that human and other mammalian immune systems can utilize different antibody gene segments to arrive at similar modes of antigen recognition, underscoring the flexibility of mammalian antibody repertoires and suggesting that experimental animals can be used to generate therapeutically-useful cross-reactive anti-RBD mAbs. The combination of mosaic-8b RBD-nanoparticles to focus the immune response and a multiplexed assay to select cross-reactive mAbs can be applied at larger scale, or against other pathogens, to identify mAbs of therapeutic and scientific potential.

## Introduction

Monoclonal antibodies (mAbs) that neutralize SARS-CoV-2 (SARS-2) have been used therapeutically to protect immunocompromised individuals and treat severe COVID-19 (1, 2). mAbs licensed for use in humans target the receptor-binding domain (RBD) of the viral spike trimer, the primary target of neutralizing Abs (3-13). With the emergence of SARS-2 variants of concern (VOCs), including the heavily mutated Omicron VOCs (14-19), previously approved therapeutic mAbs show greatly reduced or completely abrogated neutralization potencies, resulting in withdrawal from clinical use (1). Currently, only one anti-SARS-2 mAb, Pemgarda, is authorized by the FDA for humans, having been issued Emergency Use Authorization (EUA) in March 2024 (20). Pemgarda and previous anti-spike mAbs were derived from Abs isolated from SARS-2 or SARS-CoV (SARS-1) infected human donors, in some cases after in vitro selection of variants that neutralize recent VOCs (1).

The degree of cross-reactive versus variant-specific properties of an anti-RBD Ab can be understood in the context of structural studies of coronavirus spike trimers and their interactions with Abs and the SARS-2 host receptor ACE2. After binding ACE2, spike induces fusion of the viral and host cell membranes after one or more of the spike RBDs adopt an “up” position that exposes the immunodominant receptor-binding motif (RBM) to allow interactions with ACE2. Many of the most potent neutralizing Abs recognize the RBM, thereby blocking ACE2 binding (4-8, 11, 12, 21-25). We used Fab-spike structures to define classes of neutralizing anti-RBD Abs (class 1, 2, 3, 4, and 1/4) based on their epitopes, overlap with the RBM, and recognition of up and/or down RBDs on spike trimers (22, 26-28). Notably, potent class 1 and class 2 anti-RBD Abs (e.g., previously licensed therapeutic mAbs (1)), recognize epitopes that overlap with the ACE2 binding footprint, a region that exhibits high sequence variability among sarbecoviruses and early SARS-2 VOCs (22). By contrast, the epitopes of class 1/4, 4, the more recently-described class 5 (29, 30), and some class 3 Abs map to more conserved, but less accessible (in the case of class 1/4, 4, and 5 Abs), regions of sarbecovirus RBDs (22).

Targeting of conserved RBD regions could result in mAbs and vaccines that reduce the need for frequent updating. To preferentially elicit Abs against conserved epitopes, we developed mosaic-8b RBD nanoparticles (NPs) (*SI Appendix*, Fig. S1A), which display RBDs from eight different sarbecoviruses on 60-mer mi3 NPs, with the goal of stimulating B cell receptors that can crosslink using both antigen-binding Fabs between adjacent, non-identical, RBDs (31-33) (*SI Appendix*, Fig. S1B). We evaluated matched and mismatched immune responses (against viruses whose RBDs were or were not represented by an RBD on the NP), finding that mosaic-8b NPs showed enhanced heterologous binding, neutralization, and protection from sarbecovirus challenges compared with homotypic (SARS-2 RBD only) NPs in animal models (32, 34). Epitope mapping of polyclonal antisera elicited by mosaic-8b versus homotypic RBD-NPs using deep mutational scanning (DMS) (35) showed that mosaic-8b antisera primarily targeted more conserved class 4 and class 1/4 RBD epitopes that contact other portions of the spike trimer in down RBDs, whereas homotypic antiserum Abs mainly targeted variable, more accessible class 1 and 2 RBD epitopes that are not involved in intra-spike contacts (32).

Here, we describe immunization with mosaic-8b RBD-NPs as an effective strategy for identification of cross-reactive anti-sarbecovirus mAbs with therapeutic potential. We used single-cell optofluidics and multiplexed antigen specificity assays to profile antigen reactivity and assess the cross-reactivity of IgGs elicited by mosaic-8b in rabbits, finding a higher percentage of cross-reactive B cells for mosaic-8b compared with homotypic SARS-2 immunization (Fig. 1). We also mapped elicited mAb epitopes using competition ELISAs, DMS (35), 3D structure determinations, and *in vitro* selection (36, 37) and evaluated neutralization against SARS-2 variants and other sarbecoviruses (Fig. 2-7), demonstrating that the mosaic RBD-NP vaccine approach works as designed to target conserved epitopes. Thus, when paired with a multiplexed antigen specificity assay to select for cross-reactivity, mosaic RBD-NP immunization could be used to efficiently develop therapeutic neutralizing mAbs that would not be affected by current or future SARS-2 VOC substitutions and would likely remain efficacious against a future sarbecovirus spillover.

**Figure 1.**
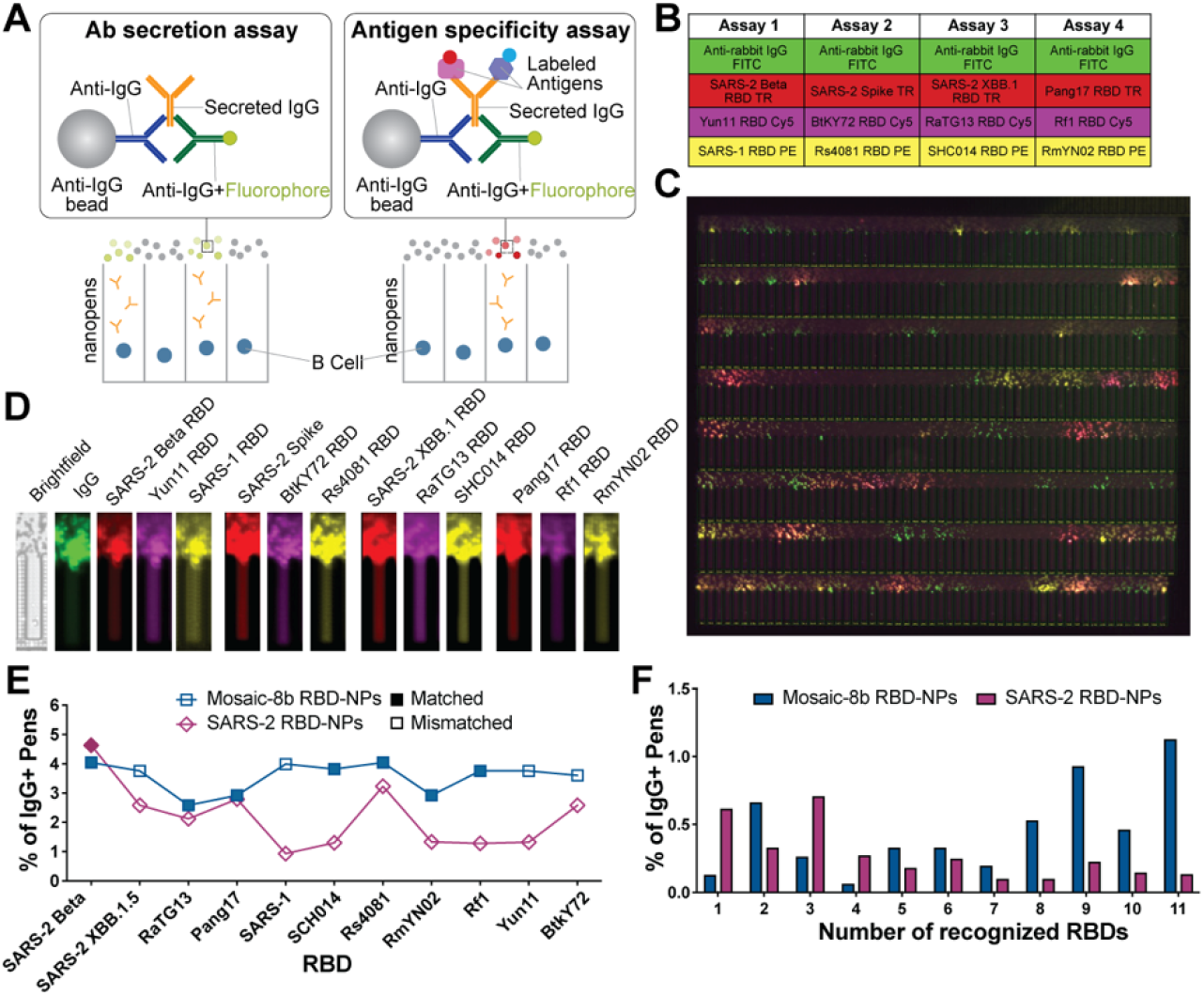
Rabbit memory B cells were characterized for secretion of broadly cross-reactive IgGs. Abbreviations. FITC: Fluorescein Isothiocyanate; TR: Texas Red; Cy5: Cyanine5; PE: Phycoerythrin. (A) Multiplexed assay to assess IgG binding breadth. Abs secreted from individual B cells in nanopens were captured by anti-rabbit IgG beads and detected with an anti-rabbit IgG secondary Ab conjugated to FITC or with antigens labeled with different fluorophores. (B) Four sequential assays for B cell screening, each of which assessed IgG secretion (FITC) and binding to three different antigens (labeled with TR, Cy5, or PE). (C) A field of view image from a multiplexed binding assay showing “blooms” above nanopens containing B cells secreting IgGs with different binding properties. (D) Individual nanopens were evaluated for antigen binding based on the presence of a bloom in fluorescent channels above the nanopen in sequential assays. Nanopens scored as positive for broad antigen binding were exported for cloning. (E) The percentage of IgG+ nanopens containing a single B cell that bind to each RBD in single-cell multiplexed assays comparing B cells from rabbits immunized with mosaic-8b RBD-NP (blue) or SARS-2 RBD-NP (mulberry). (F) The percentage of IgG+ nanopens that contain secreted IgG that recognize 1 to 11 RBDs from rabbits immunized with mosaic-8b RBD-NP (blue) or SARS-2 RBD-NP (mulberry).

**Figure 2.**
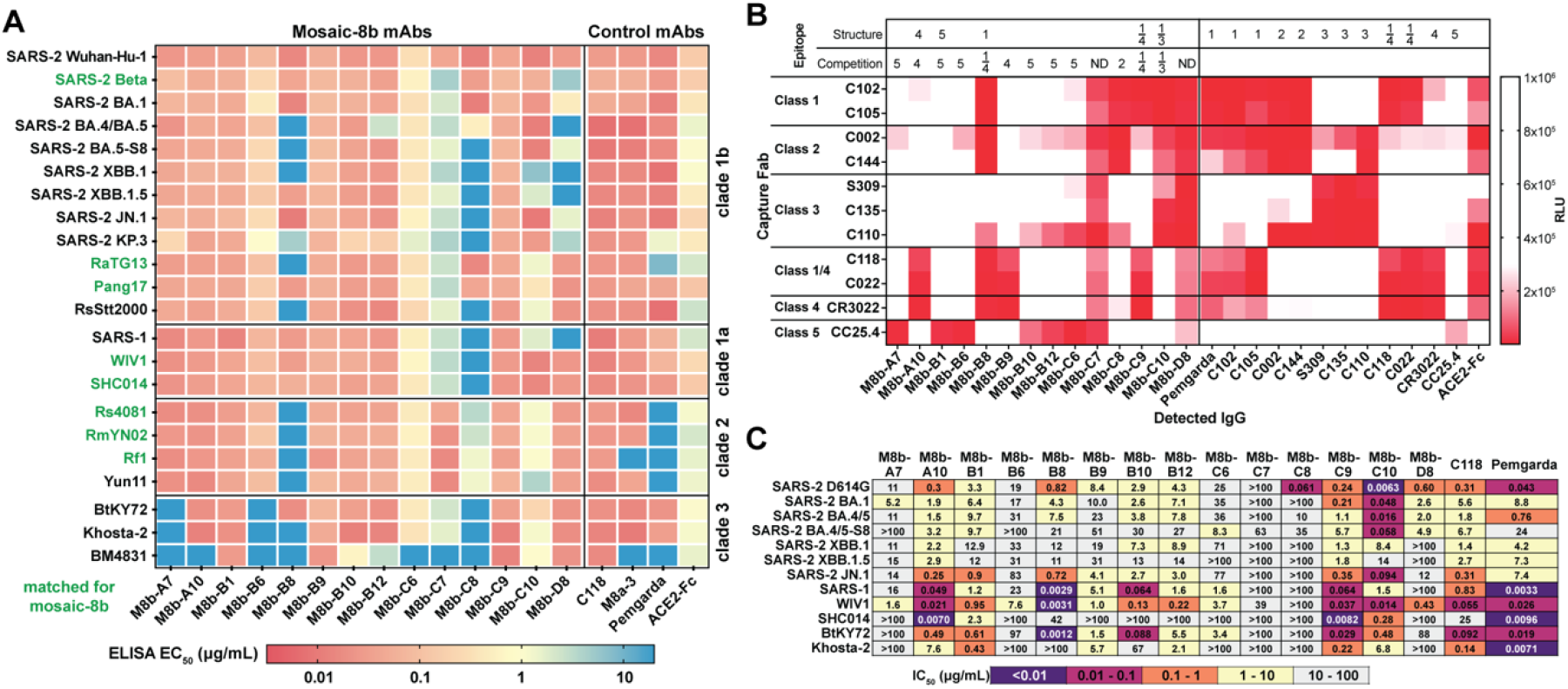
mAbs isolated from rabbits immunized with mosaic-8b exhibit broad recognition against SARS-2 and other sarbecoviruses. (A) ELISAs to assess breadth of RBD binding of mAbs isolated from rabbits immunized with mosaic-8b RBD-NPs. (B) Competition ELISAs used to identify mAb epitopes. Immobilized Fab was used to capture SARS-2 RBD, IgGs indicated at the bottom were added, and Fc was detected. White: high luminescence signal indicating IgG binding does not compete with the Fab. Red: low signal indicating reduced binding consistent with competition. The RBD epitope class for each mAb is shown based on structural analysis (Structure) or competition ELISA (Competition). ND: Not able to be determined. (C) Neutralization of rabbit mAbs against a panel of 12 pseudoviruses including SARS-2 D614G, five SARS-2 VOCs, three non-SARS-2 sarbecoviruses, and two pseudoviruses with chimeric spikes (BtKY72 or Khosta-2 RBD plus remainder of the SARS-1 spike).

## Results

### Mosaic-8b RBD-NPs elicit cross-reactive Abs

We covalently attached eight different sarbecovirus RBDs or only SARS-2 RBD to a SpyCatcher-mi3 NP (38, 39) using the SpyCatcher-SpyTag system (38, 39) to create mosaic or homotypic SARS-2 RBD-NPs (32, 34) (*SI Appendix*, Fig. S1A). Mosaic-8b RBD-NPs included SARS-2 Beta RBD plus seven other sarbecovirus RBDs from clade 1a, 1b, or 2 spikes attached randomly to 60 positions on SpyCatcher-mi3 (38), whereas homotypic SARS-2 RBD-NPs included only SARS-2 Beta RBD. In addition to previously-described characterization and validation (31, 32, 34), we used SpyCatcher- and mi3-specific mAbs and strain-specific RBD mAbs (*SI Appendix*, Fig. S1C,D, Table S1) to confirm display of all 8 RBDs on mosaic-8b RBD-NPs and only SARS-2 Beta RBD on the SARS-2 RBD-NPs (*SI Appendix*, Fig. S1E).

We next utilized a protocol to derive IgG heavy and light chain sequences from memory B cells of mosaic-8b–immunized rabbits in order to assess their potential for broad and potent anti-SARS-2 activity and to investigate whether the increased cross-reactivity of mAbs elicited by mosaic-8 observed in mice (33) extends to another species. Rabbits were primed and boosted with mosaic-8b or SARS-2 RBD-NPs, and peripheral blood mononuclear cells (PBMCs) were isolated (*SI Appendix*, Fig. S1F). Memory B cells were enriched by IgG pulldown, and recovered cells were cultured to activate IgG secretion.

We used a Beacon optofluidics system to isolate and screen ∼27,000 individual activated memory B cells for secretion of IgGs that recognized diverse sarbecovirus RBDs, both from unmatched strains (not represented by an RBD on mosaic-8b) and from matched strains (represented on the mosaic-8b or homotypic SARS-2 RBD-NPs) (Fig. 1; *SI Appendix*, Fig. S2). Assays for Ab secretion and antigen specificity were used to identify nanopens (chambers within an optofluidic chip) containing single cells secreting an IgG that bound to labeled antigens (Fig. 1A). Multiplexed assays using four fluorophores enabled screening for IgG secretion and binding to 12 antigens (Fig. 1B). Signal from fluorescence channels were overlaid to determine B cell breadth (Fig. 1C). Binding of each antigen was assessed by the presence of a fluorescent bloom above a nanopen using Beacon integrated software and manual verification (Fig. 1D).

Using results from the multiplexed assays, we found that immunization with mosaic-8b resulted in a higher percentage of B cells that recognized a range of both matched and mismatched RBDs compared with homotypic SARS-2 immunization, while SARS-2 immunization resulted in a higher percentage of B cells that recognized the matched SARS-2 Beta RBD (Fig. 1E). In addition, mosaic-8b immunization resulted in a higher percentage of B cells that recognized multiple RBDs (i.e., broadly cross-reactive mAbs) compared with homotypic SARS-2 immunization (Fig. 1F).

From the mosaic-8b RBD-NP immunized rabbit sample, we exported 90 B cells secreting IgGs that bound to any of the 12 assayed antigens, first exporting cells secreting IgGs that bound >6, followed by cells making IgGs that bound fewer antigens. The variable heavy and variable light chain genes from the first 48 exported cells were amplified and sequenced. We isolated paired heavy and light chain genes for 14 RBD-binding mAbs and subcloned them into expression vectors encoding human IgG C_H_1-C_H_2-C_H_3 domains and human C_L_ domains (for IgGs) or human C_H_1 and human C_L_ domains (for Fabs) (*SI Appendix*, Fig. S2).

### Rabbit mAbs are broadly cross-reactive

RBD recognition by 14 rabbit mAbs elicited by mosaic-8b RBD-NPs was assessed via ELISA to a panel of RBDs that included sarbecovirus RBDs from clade 1a, 1b, 2, and 3 and SARS-2 variants (Fig. 2A). Three mAbs (M8a-B1, M8a-B9, and M8a-C9) bound to all sarbecovirus RBDs tested with EC_50_ values <0.1 µg/mL, and several others bound broadly to clade 1a, 1b, and 2 RBDs, consistent with identification of cross-reactive B cells by the single-cell multiplexed assays, which were mostly predictive of ELISA results (*SI Appendix*, Fig. S3).

We further characterized mAbs to identify epitopes using a competition ELISA. Fabs derived from characterized human mAbs that recognize class 1, 2, 3, 4, 1/4, or 5 (22, 27, 29, 30) RBD epitopes were adsorbed on an ELISA plate and then incubated with SARS-2 RBD. IgGs cloned from rabbit memory B cells, control IgGs of known epitope, or a human ACE2-Fc protein (27) were added and assessed for binding (Fig. 2B). In this assay, an IgG or ACE2-Fc will only bind if it is not sterically occluded by the bound Fab. We identified likely epitopes of 12 mAbs on SARS-2 RBD including one class 2, two class 4, six class 5, one class 1/3 (competes with both class 1 and class 3 mAbs), and two class 1/4 anti-RBD mAbs (Fig. 2B). These results are consistent with previous findings that many class 3, 4, 1/4, and 5 anti-RBD Abs exhibit breadth of binding to sarbecovirus RBDs (22, 23, 27, 29, 30, 32-34, 40).

We next evaluated neutralization potencies (as assessed by inhibitory concentrations at 50%, IC_50_ values) and breadth of the rabbit mAbs against a panel of SARS-2 VOCs and other sarbecoviruses using a pseudovirus neutralization assay that correlates with authentic virus neutralization (41). Neutralization results (Fig. 2C) for particular mAbs were generally consistent with binding data (Fig. 2A). While the potencies of some mAbs were modest, M8b-A10 (class 4) and M8b-C9 (class 1/4) exhibited both potency and breadth across the panel of tested viruses (Fig. 2C). Our neutralization panel did not include pseudoviruses for clade 2 sarbecoviruses with unknown host receptors. However, ELISA binding showed that both M8b-A10 and M8b-C9 bound to the RBDs of clade 2 spikes, whereas Pemgarda showed weak or no binding to clade 2 RBDs (Fig. 2A), suggesting that this licensed therapeutic mAb would not be effective against a clade 2 sarbecovirus spillover.

### DMS reveals multiple RBD epitopes

To further evaluate residues within conserved and variable RBD epitopes (Fig. 3A) targeted by the rabbit mAbs, we used DMS to map residues that affect recognition by five mAbs (M8b-A10, M8b-B1, M8b-B8, M8b-C9, and M8b-C10) using yeast display libraries derived from SARS-2 Beta RBD (42). DMS revealed class 4 anti-RBD Ab epitope escape profiles with sensitivities to substitutions at RBD residues 378 and 411 (M8b-A10) and 411, 413, and 427 (M8b-C9) (Fig. 3B), consistent with their broad binding and neutralization (Fig. 2A,C) and competition ELISA classifications (Fig. 2B). DMS analysis of M8b-B1 showed that most escapes were mediated by substitutions that encoded a potential N-linked glycosylation site (PNGS) to add an N-glycan at RBD residue 357 (R357N) or 394 (Y396T) within the class 5 epitope (Fig. 3B), consistent with competition ELISA results (Fig. 2B) and M8b-B1’s broad binding and neutralization properties (Fig. 2A,C). DMS for M8b-B8 indicated a class 1 anti-RBD Ab profile centered on RBD residue 504 (Fig. 3B). Finally, M8b-C10 showed a class 3 profile centered on residues 441, 499, and 500, and Pemgarda showed a class 1/3 profile centered on residues 408, 500, 503, and 504, consistent with competition ELISA epitope class assignments (Fig. 2B).

**Figure 3.**
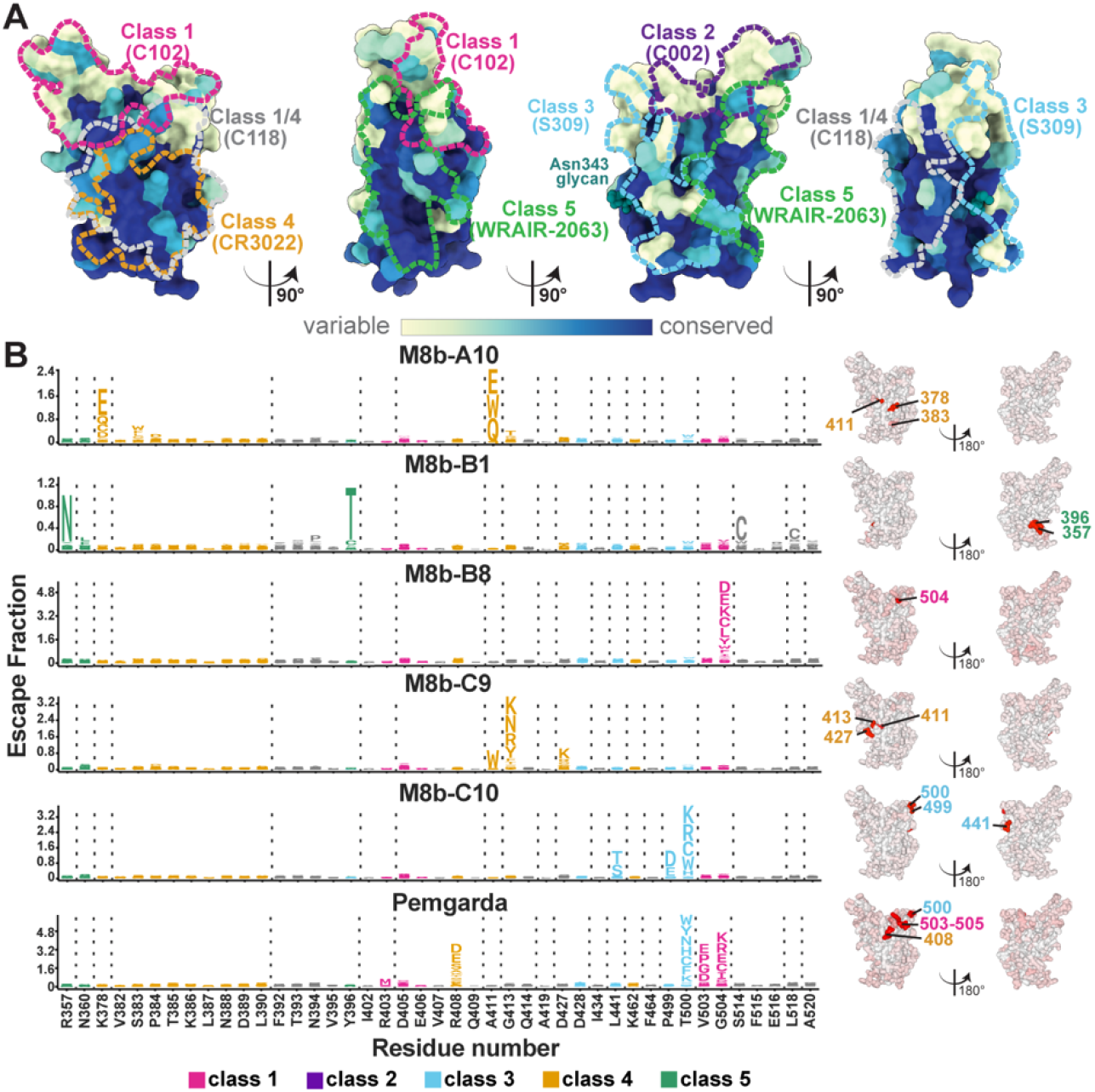
DMS of rabbit mAbs reveals RBD epitopes. (A) Sequence conservation of 16 sarbecovirus RBDs calculated using ConSurf (81) and plotted on a surface representation of SARS-2 RBD (PDB 7BZ5). Epitopes for anti-RBD class 1, 2, 3, 4, 1/4, and 5 Ab epitopes (22, 27, 29, 30) are outlined in dots in different colors using information from representative structures of mAbs bound to SARS-2 spike or RBD (C102: PDB 7K8M; C002: PDB 7K8T, S309: PDB 7JX3; CR3022: PDB 7LOP; C118: PDB 7RKV; WRAIR-2063: PDB 8EOO). (B) Results for experiments involving SARS-2 Beta yeast libraries. Left: Logo plot with RBD residue number (x-axis) and a stack of letters at RBD positions indicating amino acid substitutions that result in escape. The height of each letter indicates the degree of contribution for escape resulting from a particular substitution. The height of a stack of letters indicates the overall site-wise escape metric (as defined in ref. (74)). Letters are colored based on RBD epitope class as indicated in panel a. Right: The overall site-total escape (as defined in ref. (74)) mapped to the surface of a SARS-2 RBD (PDB 6M0J), with white indicating no escape and red indicating escape. Lines with colored numbers according to epitope class indicate RBD positions with the most escape.

### 3D structures rationalize mAb properties

To further explore recognition and neutralization mechanisms, we solved 3.2 Å-3.5 Å resolution single-particle cryo-electron microscopy (cryo-EM) structures of rabbit mAb Fabs complexed with a SARS-2 spike-6P trimer (43) and 2.4 Å-2.6 Å resolution X-ray structures of Fab-RBD complexes (Fig. 4,5; *SI Appendix*, Fig. S4, Tables S2,3). As previously observed for structures of infection- and vaccine-induced human mAb Fabs bound to SARS-2 spike trimers (22, 25, 44, 45), the rabbit mAb Fab cryo-EM structures showed different modes of RBD recognition: M8b-A10, M8b-B8, and M8b-C9 Fabs bound to RBDs only when they were in an “up” conformation (Fig. 4A-C), and M8b-C10 Fab recognized both “up” and “down” RBDs (Fig. 4D). Because only dissociated trimers were observed on cryo-EM grids of M8b-B1 Fab incubated with a spike trimer, we determined a crystal structure of a M8b-B1 Fab-RBD complex (Fig. 4E).

**Figure 4.**
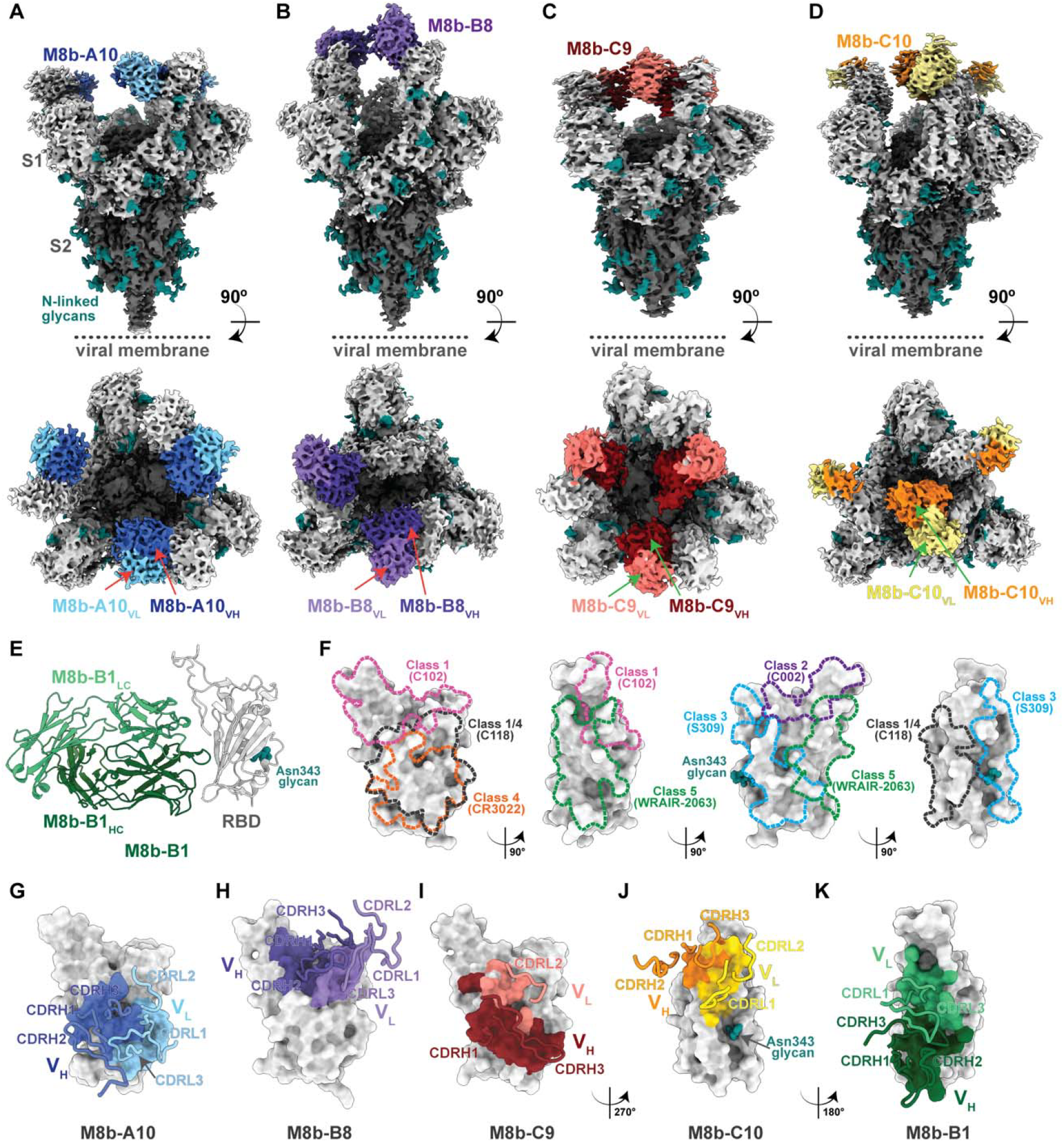
Rabbit mAbs recognize various epitopes on SARS-2 RBDs. (A-D) Side and top-down views of EM densities of cryo-EM structures of Fab-spike complexes (V_H_-V_L_ densities shown for each Fab) for (A) M8b-A10, (B) M8b-B8, (C) M8b-C9, and (D) M8b-C10. (E) Crystal structure of a M8b-B1 Fab-RBD complex. (F) Epitopes of representative human mAbs from five anti-RBD Ab classes outlined in colored dots using structural information (C102: PDB 7K8M; C002: PDB 7K8T, S309: PDB 7JX3; CR3022: PDB 7LOP; C118: PDB 7RKV; WRAIR-2063: PDB 8EOO). (G-K) Rabbit mAb epitopes with interacting CDRs on a surface representation of SARS-2 RBD (gray) shown as footprints for the indicated V_H_ and V_L_ domains for epitopes with superimposed CDR loops for those CDRs that contact the RBD. CDRs were assigned using Kabat definitions (82).

**Figure 5.**
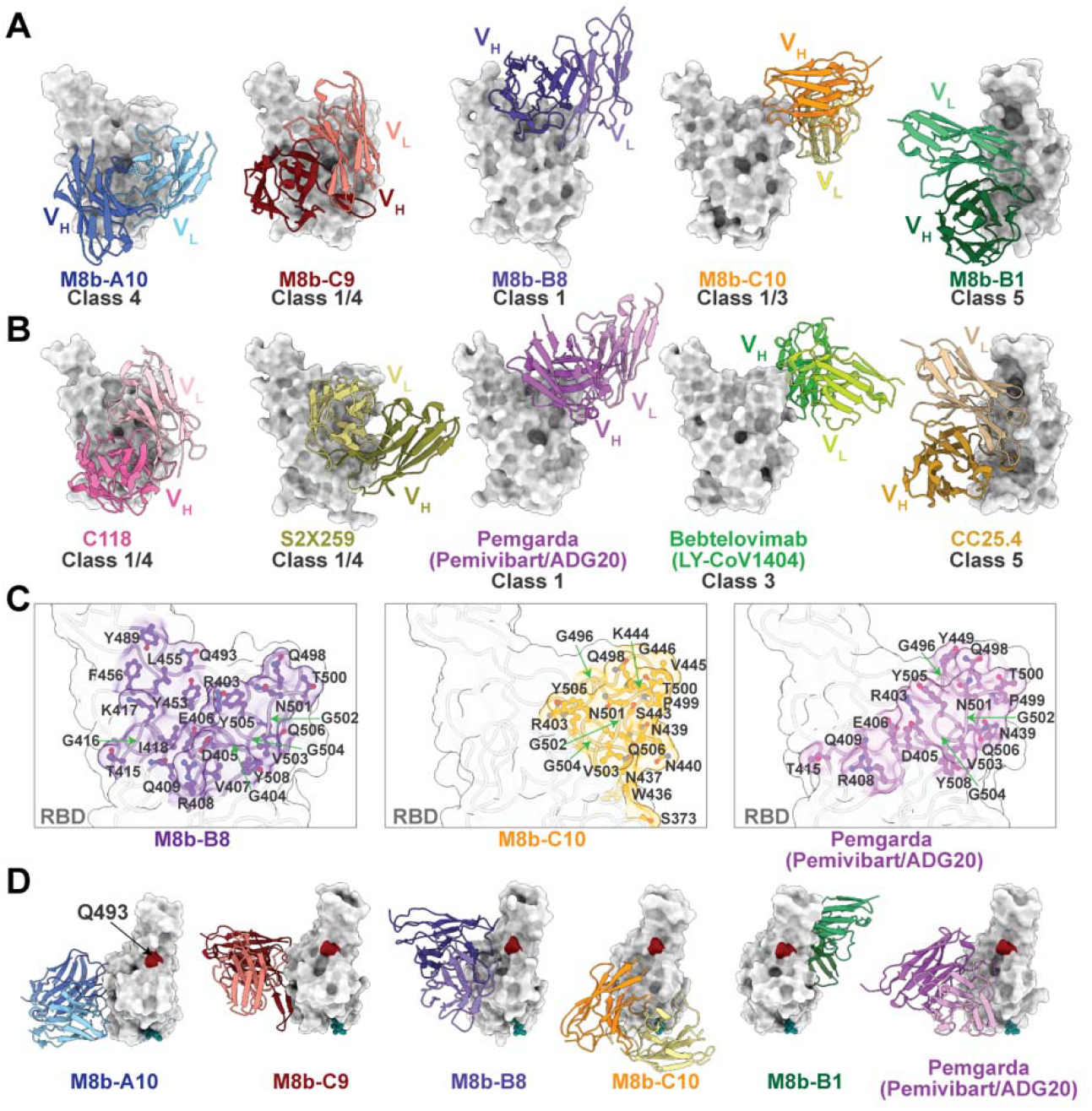
Rabbit and human mAbs recognize similar epitopes. (A,B) V_H_-V_L_ domains of mAbs (cartoon representations) (rabbit, panel A; human, panel B) complexed with SARS-2 RBD (surface representations). Human anti-RBD mAbs (C118: PDB 7RKV; S2×259: PDB 7M7W; Pemgarda (Pemivibart/ADG20): PDB 7U2D; Bebtelovimab (LY-CoV1404): PDB 7MMO; WRAIR-2063: PDB 8EOO) with similar epitopes are shown below each rabbit mAb-RBD complex. (C) Comparisons of RBD epitopes of selected rabbit mAbs with Pemgarda. RBD epitope residues (determined by PDBePISA (83)) shown as sticks. (D) Comparison of RBD recognition by rabbit mAbs and Pemgarda. The position of RBD residue Q493, which is substituted in recent VOCs that show resistance to Pemgarda (49), is highlighted in dark red.

The M8b-A10–spike cryo-EM structure revealed that each of the three “up” RBDs was recognized by an M8b-A10 Fab using all six of its complementarity-determining regions (heavy chain CDRH1, CDRH2, and CDRH3 and light chain CDRL1, CDRL2, and CDRL3) and heavy chain framework region 1 (FWRH1) (Fig. 4A,G, *SI Appendix*, Fig. S5A). Consistent with competition ELISA and DMS results (Fig. 2B, 3B), the footprint of M8b-A10 on the RBD surface resembled that of a cross-reactive class 4 anti-RBD mAb, CR3022 (40) (Fig. 4F,G), which was originally isolated from a SARS-1–infected patient (46). Also consistent with M8b-A10’s cross-reactive binding and neutralization properties (Fig. 2A,C), its binding footprint involved residues that are largely conserved across sarbecovirus RBDs (Fig. 3A,4G).

The M8b-B8 Fab-spike structure showed two Fabs binding to “up” RBDs with the third, non-Fab-bound, RBD adopting the “down” position (Fig. 4B). RBD recognition was mediated by both the V_H_ and V_L_ domains using all six CDRs plus FWRH1 (Fig. 4H, *SI Appendix*, Fig. S5B). We classified M8b-RBD as a class 1 anti-RBD Ab because the majority of RBD residues at the M8b-B8 epitope were residues from the variable class 1 epitope (RBD residues 403, 405, 415-417, 453, 455, 456, 489, 493, 498, 500-503, and 505), although two RBD residues (Y453 and Y489) overlapped with the variable class 2 RBD epitope and one residue (R408) overlapped with the more conserved class 4 RBD epitope (Fig. 3A, 4F,H, *SI Appendix*, Fig. S6A-C). This is consistent with competition ELISA results showing that M8b-B8 competed with mAbs from class 1, class 2, class 4, and class 1/4 mAbs (Fig. 2B). In addition, although their binding epitopes do not overlap, M8b-B8 competed with C110, a human class 3 anti-RBD mAb (22), likely due to steric clashes between the C_H_1 and C_L_ domains of the bound Fabs (*SI Appendix*, Fig. S6D). Binding to non-conserved regions of the RBD contributed to limited binding of M8b-B8 to sarbecovirus RBDs by ELISA and weak neutralization potencies against a subset of sarbecoviruses and variants of SARS-2 (Fig. 2A,C).

We observed three M8b-C9 Fabs bound to spike in the cryo-EM structure, all recognizing “up” RBDs (Fig. 4C), as found for M8b-A10, which also recognized only “up” RBDs (Fig. 4A). To resolve high-resolution details of the Fab-RBD interaction, we solved a 2.6 Å crystal structure of an M8b-C9–SARS-2 RBD complex (*SI Appendix*, Table S3). Recognition of M8b-C9’s class 1/4 RBD epitope was mediated mainly by its V_H_ domain (Fig. 4I) through heavy chain CDRH1, CDRH3, FWRH1, and FWRH2 and light chain CDRL2 and FWRL2 regions (*SI Appendix*, Fig. S5C). The identification of a class 1/4 epitope in the Fab-spike and Fab-RBD structures is consistent with competition for RBD binding with class 1 and class 4 anti-RBD mAbs (Fig. 2B), cross-reactive binding of M8b-C9 IgG to all spike proteins and RBDs (Fig. 2A), and broad neutralization of SARS-2 variants and other sarbecoviruses (Fig. 2C).

The M8b-C10 Fab-spike structure revealed Fabs binding to all spike RBDs, with two RBDs adopting an “up” conformation and the third RBD in the “down” conformation (Fig. 4D). The interaction between M8b-C10 and the RBD was mediated by its heavy chain CDRH1, CDRH2, CDRH3, and FWRH1 and light chain CDRL1 and CDRL2 (Fig. 4J, *SI Appendix*, Fig. S5D). As seen by comparing the outlines of Ab epitopes on an RBD surface (Fig. 4F), the M8b-C10 binding epitope overlaps with the epitopes of class 1 and class 3 anti-RBD mAbs, but not with epitopes of class 2 or class 4 mAbs (*SI Appendix*, Fig. S6E). This class 1/3 epitope (Fig. 2B) mostly involves residues that vary between sarbecoviruses (Fig. 3A), consistent with the limited RBD recognition observed by ELISA for M8b-C10 (Fig. 2A). Although the binding footprints of M8b-C10 and the class 2 mAb C002 do not overlap (*SI Appendix*, Fig. S 6E), M8b-C10 competed with two class 2 mAbs for RBD binding (Fig. 2B), most likely due to steric clashes between Fab C_H_1 and C_L_ domains.

The M8b-B1 Fab-RBD crystal structure showed recognition of an epitope on a side of RBD that is buried in “down” RBDs (Fig. 4E). The interaction between M8b-B1 and RBD was mediated by FWRH1 and all CDR loops except CDRL2 (Fig. 4K, *SI Appendix*, Fig. S5E). Modeling suggested that M8b-B1 Fab would clash with a neighboring NTD when recognizing a trimer with “up” RBDs (*SI Appendix*, Fig. S5F), consistent with M8b-B1 Fab incubation with spike resulting in dissociated spike trimers on cryo-EM grids. The structurally-identified M8b-B1 epitope (Fig. 4K) is consistent with competition with CC25.4 (Fig. 2B), which recognizes the partially conserved class 5 RBD epitope (29, 30), and M8b-B1 cross-reactive binding and potent neutralization against most SARS-2 variants and other sarbecoviruses (Fig. 2A,C).

We also compared structures of the rabbit mAbs to structures of mAbs that were isolated from COVID-19 convalescent donors and FDA-approved mAbs (Fig. 5). M8b-A10 and M8b-C9 targeted similar regions of the RBDs as C118 (27) and S2×259 (47), two cross-reactive and potent mAbs that were isolated from convalescent human donors (Fig. 5A,B). Importantly, M8b-B8 and M8b-C10 both recognize epitopes similar to that of Pemgarda, the only currently FDA-approved therapeutic mAb for SARS-2 treatment (20), and Bebtelovimab, a previously FDA-approved anti-SARS-2 mAb (1) (Fig. 5A,B). M8b-B8 and Pemgarda share common epitope residues, including RBD residues 403, 405, 406, 408, 409, 415, 498, 500-505, and 508, and M8b-C10 and Pemgarda share RBD epitope residues 403, 439, 496, and 498-506. Although reside Q493 was not classified by PDBePISA (48) as part of the binding epitope of Pemgarda (Fig. 5C), a Q493E substitution was shown to contribute to Ab evasion of Pemgarda (49). Only one of the rabbit mAbs (M8b-B8) targets an RBD epitope containing residue Q493 (Fig. 5D, *SI Appendix*, Fig. S5B).

### Cross-species class 1/4 mAb features

Identification of rabbit (and previously mouse (33)) anti-RBD mAbs raised by mosaic-8 immunization offers the opportunity to compare their properties with those of human mAbs raised by SARS-2 infection and or COVID-19 vaccination; e.g., since V, D, and J gene segments can differ between species (50), it is of interest to ask whether recognition features are conserved in anti-RBD mAbs raised in different species.

Of relevance to this question, a public class of anti-SARS-2 human mAbs with a CDRH3 YYDxxG motif derived from the D gene segment of IGHD3-22 was identified (51, 52) (Fig. 6). Human mAbs in this recurrent class, e.g., COVA1-16 (26), C022 (27), and ADI-622113 (51) use the YYDxxG motif to recognize the class 1/4 RBD epitope by extending an RBD *β*-sheet through mainchain hydrogen bonding with a two-stranded antiparallel *β*-sheet formed by the CDRH3 (Fig. 6A-C), as originally described for a COVA1-16–RBD structure (26). Structural comparisons of the interfaces of RBD with M8b-C9 (Fig. 6F) and ADI-62113 (53) (Fig. 6C) revealed that heavy chain residues Y98_VH_ and Y99_VH_ in M8b-C9 interact with RBD with the same interface geometry as the tyrosines in the YYDxxG motif. A structural motif search for Abs with a YY motif in CDRH3 that interacts with RBD residues 378-382 found other examples: human mAbs CC25.36 (53) (Fig. 6D) and S2×259 (47) (Fig. 6E), as well as several nanobodies (54). CC25.36 was also previously noted to use a CDRH3 YYDML motif to recognize RBD in a manner similar to YYDxxG Abs (53). The genetic origins of the core YY features within CDRH3s vary: IGHD3-22 for YYDxxG motif Abs, IGHD3-9 for YYDML motif Abs, and a CDRH3 random library for the synthetic nanobodies. IMGT/V-Quest analysis (55) of the M8b-C9 V_H_ gene segment revealed that its YY motif derived from N region addition rather than from a D gene segment. The smallest core motif for the above Abs could be considered to be the tyrosine (Y98_VH_ in M8b-C9) that interacts with the aliphatic portion of the RBD K378 sidechain in combination with adjacent backbone hydrogen bonds to RBD residues 378-379 that extend an antiparallel *β*-sheet of the RBD. This feature was described for mAb C118 (27), which contains a YT CDRH3 sequence (Fig. 6G), and the structural motif search found additional cases: Ab AB-3467 (28) (YS) (Fig. 6H) and Ab N3-1 (56) (YF) (Fig. 6I). These comparisons provide examples in which the immune system converges upon similar recognition motifs derived either from different Ab gene segments or from processes such as N-region addition that occur during V-D-J recombination.

**Figure 6.**
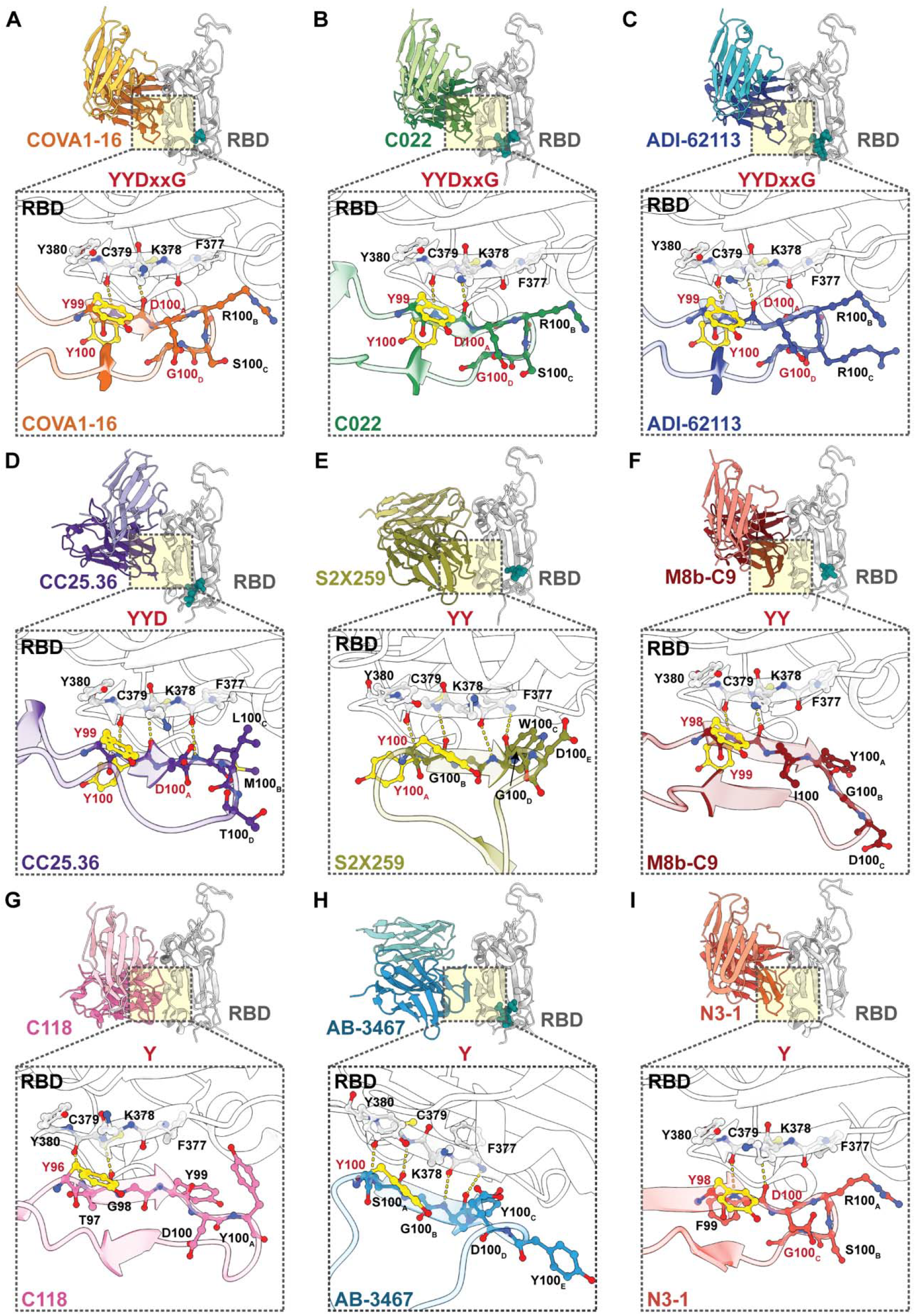
Comparisons of YYDxxG-, YY-, and Y-containing CDRH3 loops in class 1/4 anti-RBD mAbs. V_H_-V_L_–RBD complexes (cartoon diagrams) shown for indicated mAbs with zoomed-in insets (yellow shading) showing interactions between mAb CDRH3 loops and RBD as sticks, mainchain hydrogen bonds between a CDRH3 and RBD residues 377-380 as yellow dots, and the sidechains of YY and Y residues within CDRH3 motifs in yellow ball-and-stick representation. RBD (gray) complexes shown for (A) COVA1-16 (PDB 7S5R), (B) C022 (PDB 7RKU), (C) ADI-62113 (PDB 7T7B), (D) CC25.36 (PDB 8SIQ), (E) S2×259 (PDB 7M7W), (F) M8b-C9, (G) C118 (PDB 7RKV), (H) AB-3467 (PDB 7MSQ), and (I) N3-1 (PDB 8TM1). mAb-RBD figures are grouped into YYDxxG, YY, and Y motifs, with names of residues in CDRH3 motifs in red.

### Selection reveals functional escapes

To investigate the sensitivity of the rabbit mAbs to viral escape, we conducted cell culture selection experiments (57, 58) using a replication-competent recombinant vesicular stomatitis virus (rVSV) encoding the spike protein of either SARS-2 Wuhan-Hu-1, SARS-2 BA.2, SARS-2 XBB.1.5, or SARS-1 (41). For each experiment, the corresponding rVSV was allowed to replicate in the presence of a rabbit Ab at a concentration 10-fold above its IC_50_ value (Fig. 2C) to select for escape variants (Fig. 7). Consistent with structural results (Fig. 4A,G), M8b-A10 selected for escape at residues within the class 4 RBD epitope of both SARS-2 and SARS-1 spike-bearing viruses (Fig. 7A). Escape from M8b-A10 included substitutions at K378 (92.7% K378Q and 6.6% K378E in the Wuhan-Hu-1 chimeric virus), an RBD position that was identified as a key residue of escape using DMS (Fig. 3B), and at G413 (26.6% and 29.2% G413R in the BA.2 and XBB.1.5 contexts, respectively) and the equivalent residue in SARS-1, G400R (94.9% substitution). Selection in the presence of the class 5 mAb M8b-B1 resulted in viruses containing substitutions in the RBD outside of the mAb epitope, including P384L (46.6% substitution in Wuhan-Hu-1), as well as within the epitope (E465K, 59.3% substitution in XBB.1.5 and E452G or E452D, 97.9% substitution in SARS-1 viruses) (Fig. 7B). In contrast, DMS showed selection by addition of PNGSs within the epitope, substitutions that have not been seen to date in viral isolates (59), perhaps because N-glycan addition in the context of a replicating virus in viral isolates and in this selection system (57, 58) could reduce viral fitness or infectivity. Escape from M8b-C9, a class 1/4 mAb, led to the acquisition of a G413R substitution in SARS-2 spikes (98.7% in Wuhan-Hu-1, 30.7% in BA.2, 98.6% in XBB.1.5) and D414V in SARS-1 (78.0%) (Fig. 7C). These RBD residues constitute some of the more variable positions within the conserved class 4 RBD epitope (Fig. 3A). Selection of SARS-2 chimeric viruses in the presence of M8b-C10, a class 1/3 mAb whose RBD binding footprint overlaps with that of Pemgarda (Fig. 5C), resulted in a virus with a P499Q substitution in Wuhan-Hu-1 RBD (100%) (Fig. 7D, which was also identified as a site of escape using DMS (Fig. 4B). Escape pathways of the BA.2-bearing virus were more diverse, with substitutions observed at 8 positions in and around the epitope, including residues in class 1, 3, and 4 RBD epitopes. Three substitutions observed during BA.2 selection in the presence of M8b-C10 were reversions to the Wuhan-Hu-1 sequence (A376T, N405D, S408R), and two additional substitutions are found in variant lineages (e.g., G446S in BA.1, XBB.1.5, KP.3; G496S in BA.1). With the exception of BA.2 selection in the presence of M8b-C10, all other substitutions observed in these selection experiments are seen only at low frequencies in natural sequences (<0.03%) (59).

**Figure 7.**
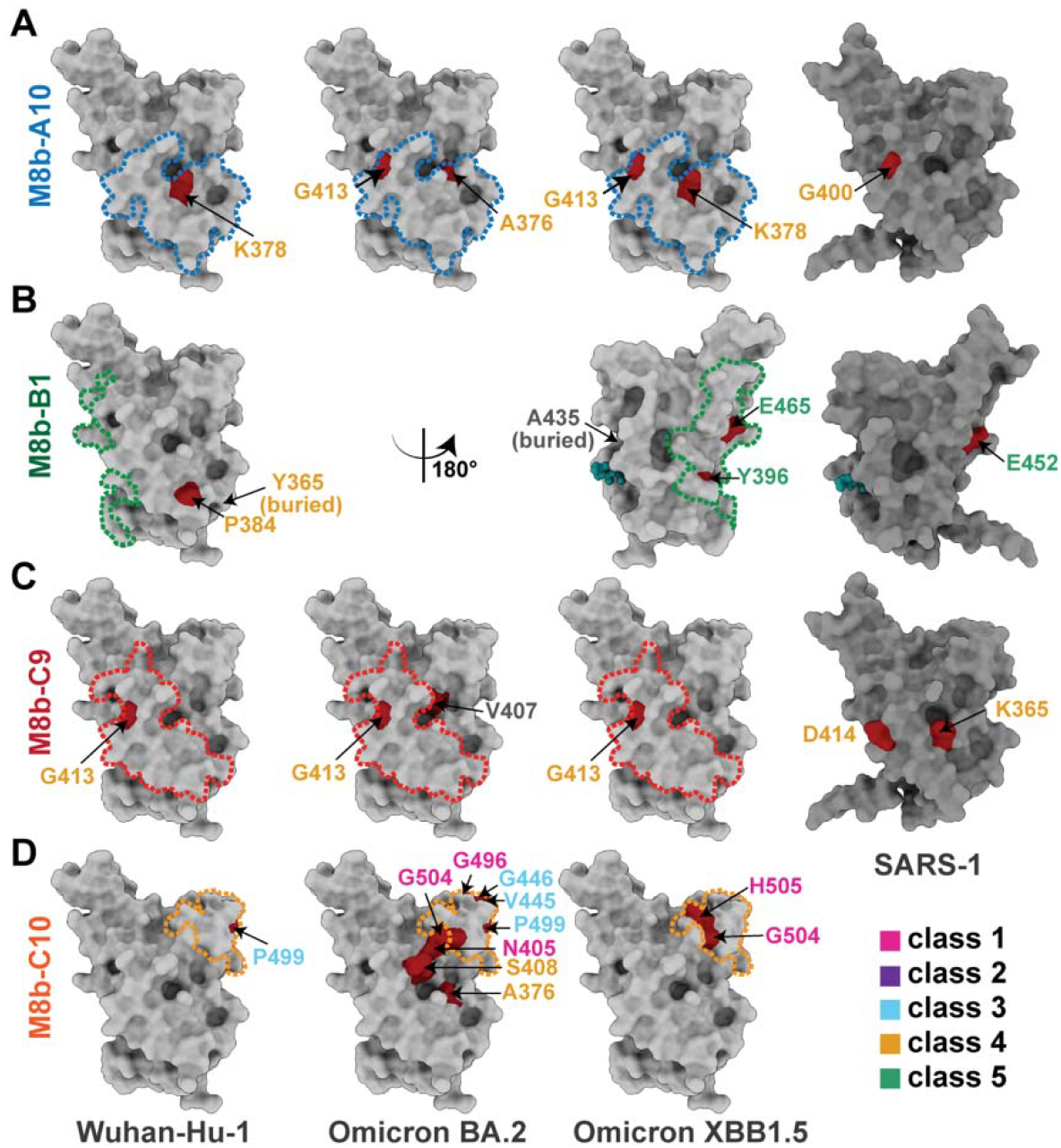
Selection experiments reveal functional escapes. Positions of residues (highlighted in red on RBD surfaces) in SARS-2 and SARS-1 RBD that were substituted following passage of rVSVs encoding the spike proteins of SARS-2 Wuhan-Hu-1, Omicron BA.2, Omicron XBB.1.5, or SARS-1 with (A) M8b-A10, (B) M8b-B1, (C) M8b-C9, or (D) M8b-C10. Residue names are labeled in colors according to RBD epitope class. Epitopes for these mAbs are outlined in dots colored according to epitope class.

## Discussion

Therapeutic mAbs against the SARS-2 spike were beneficial for many during the COVID-19 pandemic (1, 60). However, emergence of previous, and likely future, VOCs with mutations that reduce mAb neutralizing activities against variants, along with recommendations against the use of single mAbs therapeutically (61), provide a compelling rationale to develop processes by which additional mAbs can be efficiently generated and identified, especially mAbs that broadly recognize RBDs from SARS-2 VOCs and animal sarbecoviruses.

As of 2024, the only approved mAb for COVID-19 treatment is Pemgarda, which is authorized for pre-exposure prophylaxis as a treatment option for immunocompromised individuals who do not make strong responses to vaccines (20). However, given that Pemgarda’s RBD epitope spans a region that exhibits variability (Fig. 3A, 5B), emergence of future VOCs are likely to limit its efficacy, consistent with a report that Pemgarda exhibits diminished neutralization of KP.3.1.1, a rapidly expanding VOC lineage (49). Most therapeutic anti-SARS-2 mAbs have been isolated from infected and/or immunized human donors (62), in whom immune responses generally focus on the immunodominant, but variable, class 1 and class 2 RBD regions (4-8, 11, 12, 21-25), thus usually resulting in mAbs that are sensitive to VOC substitutions. In addition, therapeutic mAbs developed against SARS-2 often required long development times; e.g., Pemgarda was engineered from a mAb originally isolated from a SARS-1 patient and then affinity matured (62, 63), necessitating a multi-year process.

Here, we suggest a more directed and efficient approach to identifying cross-reactive mAbs; namely, immunizing with mosaic-8 NPs, which we have demonstrated direct Ab responses to more conserved RBD regions (31, 32, 34, 64), and prescreening for breadth using a multiplexed binding assay. We previously reported identification of cross-reactive mAbs elicited in mice immunized with mosaic-8 NPs (33). Here, we report discovery of promising mAbs exhibiting broad neutralization across animal sarbecoviruses derived from mosaic-8b–immunized rabbits. We chose rabbits for immunization because they have diverse Ab repertoires and were previously used for isolation of potent anti-SARS-2 spike RBD mAbs (65, 66). In the current study, we used multiplexed assays to evaluate binding to multiple antigens by IgGs secreted by individual B cells for comparing the breadth of responses to mosaic-8b versus homotypic SARS-2 RBD-NPs, finding broader responses for B cells from mosaic-8b– than homotypic SARS-2– immunized animals. In a screen of only 14 mAbs, we identified M8b-B8 and M8b-C10 with epitopes that overlap with and/or resemble Pemgarda’s (Fig. 5A-C), as well as cross-reactive class 4 (M8b-A10) and class 1/4 (M8b-C9) mAbs (Fig. 2, 5A). A larger screen could be used to identify more mAbs with broadly cross-reactive neutralization profiles, which could be used therapeutically in combination. For identifying mosaic-8b–induced mAbs for therapeutic use in humans, mosaic-8b immunization could be done in transgenic human immune repertoire mice (67), wildtype animal-elicited mAbs could be humanized (68), and/or human mAbs isolated from participants in an upcoming mosaic-8b clinical trial funded by the Coalition for Epidemic Preparedness Initiative (CEPI).

Our results showed that differing Ab gene segment repertoires between humans and animals did not prevent mosaic-8b elicitation in rabbits of broadly cross-reactive Abs, including those with recognition properties similar to human anti-RBD Abs. For example, there is no apparent rabbit counterpart of the human D gene segment IGHD3-22, which encodes a common CDRH3 YYDxxG motif (52) that recognizes a class 1/4 RBD epitope by extending an RBD *β*-sheet through hydrogen bonding with an Ab CDRH3 *β*-strand. Yet with a relatively small screen, we found a rabbit mAb, M8b-C9, that uses a portion of the YYDxxG motif to extend the same RBD *β*-sheet using its CDRH3. In addition, two human mAbs, C022 and C118, exhibit RBD recognition properties resembling those of COVA1-16, yet only C022 contains the YYDxxG motif in its CDRH3 (27). Thus, the immune systems of humans and other mammals can utilize different Ab gene segments to arrive at similar modes of antigen recognition, underscoring the flexibility of mammalian Ab repertoires and suggesting that animal models can be used for screens to identify cross-reactive anti-RBD mAbs of potential therapeutic utility. It should also be possible to use a multiplexed assay, as described here, to select for broadly cross-reactive Abs from mosaic-8b– immunized humans as potential therapeutic mAbs.

## Materials and Methods

Additional Method details can be found in the *SI Appendix*.

### RBD expression and production of RBD-NPs

SpyTag003 (39)-tagged RBDs for conjugation, Avi-tagged RBDs for Beacon assays and ELISAs, and soluble SARS-2 Wuhan-Hu-1 spike trimer with 6P stabilizing mutations (43) were expressed by transient transfection in Expi293F cells and purified as described (32, 34). Human IgG mAbs, rabbit IgG mAbs, and human ACE-2 fused to human IgG Fc (hACE2-Fc) (27) were expressed and purified as described (22, 27). Fabs for ELISAs were produced by papain cleavage or expressed as His-tagged Fabs as described (33). SpyCatcher003-mi3 NPs (38) were produced in *B. subtilis* (Ingenza, LTD) and purified as described (31).

RBD-NPs were generated and purified as described (32). Concentrations of conjugated RBD-NPs are reported based on RBD content determined using a Bio-Rad Protein Assay.

### mAbs for characterization assays

Rabbit mAbs that bind to SpyCatcher (anti-SC003 E1), mi3 (anti-mi3 A3), and RaTG13 (anti-RaTG13 F1) were identified from memory B cells from rabbits immunized with mosaic-8b or homotypic RBD-NPs. Activated PBMCs were loaded on a Beacon chip and assayed for binding to SpyCatcher, mi3, or RBDs. Candidate cells were exported and VH and VL gene segments were cloned into expression vectors encoding human constant domains. Human mAbs that bind to Pang17, SHC014, Rs4081, Rf1, or RmYN02 RBD but not the other seven RBDs on mosaic-8b RBD-NPs (anti-Pang17, anti-SHC014, anti-Rs4081, Anti-Rf1 422, Anti-Rf1 425, Anti-Rf1 428, anti-RmYN02) were identified using HuCAL technology with a positive selection and multiple negative selection strategy (Biorad) and cloned into expression vectors encoding human constant domains. *β*38 IgG (binds SARS-2 Beta RBD) and M8a-7 IgG (binds WIV1 RBD) were described previously (33, 69).

### Mosaic-8b RBD-NP rabbit immunizations

Immunizations were performed by Labcorp Drug Development using IACUC-approved protocols. Five 7-8 week old New Zealand White female rabbits were immunized intramuscularly with 50 µg of mosaic-8b RBD-NPs in 50% v/v AddaVax^™^ adjuvant. One female rabbit was immunized intramuscularly with 50 µg of homotypic SARS-2 RBD-NPs in 50% v/v AddaVax^™^ adjuvant and boosted 4 weeks later. Peripheral blood mononuclear cells (PBMCs) and serum were obtained and stored at -80 °C.

### Bruker Cellular Analysis Beacon assays

Rabbit PMBCs were MACS-enriched as described by Bruker Cellular Analysis in the rabbit Memory B cell workflow and activated in Activation Media for 4 days. Activated PBMCs were loaded on the Bruker Cellular Analysis Beacon instrument and penned into individual nanopens similar to previous reports (33). Assays were multiplexed using goat anti-rabbit IgG (Fc) Coated Polystyrene Particles (Spherotech), Alexa Fluor 488 goat anti-rabbit IgG, and soluble biotinyated antigens coupled to streptavidin-fluorophore conjugates.

Nanopens were scored for IgG secretion and antigen capture and cells of interest were exported. Ig gene sequences of interest were obtained as instructed by Bruker Cellular Analysis. PCR products from a cDNA library generated from the mRNA of the exported cell were sequenced cloned into expression constructs encoding human constant domains.

### Binding and Competition ELISAs

ELISAs were performed using a Tecan Evo liquid handling robot as described (31). Where indicated, curves were plotted and integrated to obtain half-maximal effective concentrations (EC_50_) using Graphpad Prism v9.3.1 assuming a one-site binding model with a Hill coefficient. Data points represent the mean and error bars represent the standard deviation of four replicates.

Competition ELISAs were performed using a Tecan Evo liquid handling robot as described (31) using Fabs corresponding to mAb of known epitope adsorbed to the plate, followed by blocking, SARS-2 Wuhan-Hu-1 RBD or NP, then IgG. Bound IgG was detected using horseradish peroxidase-conjugated Goat Anti-Human IgG Fc. Measurements were performed in quadruplicates and means are shown in a heat map.

### Pseudovirus neutralization assays

Neutralization assays using pseudoviruses based on HIV lentiviral particles were conducted as described (8, 70). Half-maximal inhibitory concentrations (IC_50_ values) were determined using nonlinear regression in AntibodyDatabase (71).

### DMS

DMS studies were performed in duplicates using SARS-2 Beta RBD libraries (generously provided by Tyler Starr, University of Utah) as described (31, 72). Stained yeast cells were sorted to capture RBD mutants that had reduced mAb binding but relatively high RBD expression. mAb-escaped cells were expanded and DNA extraction and Illumina sequencing were carried out as described (73). Escape fractions were computed using processing steps described (73, 74) and implemented using a Swift DMS.

Logo plot visualizations of escape maps were created using Swift DMS (73), where letter height indicates the escape score for that amino acid mutation, and height of the stack of letters indicate the total site-wise escape metric, calculated as described (73). Letters for each site were colored according to epitope class. Structural visualizations were performed as described using an RBD surface (PDB 6M0J) colored by the site-wise escape metric at each site (31).

### *In vitro* selection experiments

To identify viral escape substitutions in the presence of mAb, we used a recombinant replication-competent vesicular stomatitis virus (rVSV) encoding the spike proteins of either SARS-2 Wuhan-Hu-1, SARS-2 VOC BA.2, SARS-2 VOC XBB.1.5, or SARS-1 as described (41, 57). Viral populations were incubated with mAb at a concentration 10x above its IC50 value and then added to HEK-293T/ACE2cl.22 cells. Following the second passage, RNA was extracted from filtered supernatant and reverse-transcribed. Sequences encoding the extracellular domain of spike were amplified and sequenced using Illumina MiSeq to identify mAb escape substitutions in the RBD. Sequencing reads were aligned to the corresponding RBD reference sequence and annotated for the presence of mutations. A variant was defined as occurring at a frequency >3% of reads at that position.

### Cryo-EM and X-ray crystallography

Complexes of SARS-2 Wuhan-Hu-1 spike and Fabs were prepared and frozen as described (33). Single-particle cryo-EM datasets for complexes were collected using SerialEM (75) on a 300 keV Titan Krios or a 200 keV Talos Arctica and data were processed using cryoSPARC v4.3 (76) as described (33).

Initial models were generated by docking Fab–SARS-2 RBD structures into the locally refined cryo-EM density using UCSF Chimera (77), followed by docking the remaining SARS-2 spike trimer (PDB 7SC1) into the cryo-EM density map. The model was refined in Phenix (78) using real space refinement and the amino acid sequences for the mAbs were manually corrected in Coot (79). Single-particle cryo-EM statistics are reported in *SI Appendix*, Table S2.

Crystallization trials for Fab–SARS-2 RBD complexes were set up using commercially available screens as described (33) and x-ray diffraction data were collected at the Stanford Synchrotron Radiation Lightsource (SSRL) beamline 12-2. X-ray datasets were processed, solved with molecular replacement, and refined using Phenix (78) and Coot (79) as described (33). Crystallographic statistics are reported in *SI Appendix*, Table S3.

### Structural Motif Searches

To identify Ab/RBD structures with a YY motif in CDRH3 that interacts with RBD residues 378-382, we used the structure motif search service at RCSB (80) with PDB ID 7RKU; residues A51, A52, A53, A54, A55, G104, and G105; RMSD cutoff of 2 Å; and an Atom Pairing setting of All Atoms. A second structure motif search was done for a single CDRH3 tyrosine (corresponding to Y98_VH_ in M8b-C9) and RBD residues 378-379 with PDB ID 7RKU; residues A51, A52, G104, and G105; with exchanges of G105 to all amino acids; RMSD cutoff of 1.5 Å; and an Atom Pairing setting of Backbone Atoms.

## Data availability

All data supporting the findings of this study are available within the paper and *SI Appendix*. Raw sequencing data from DMS experiments are deposited on NCBI SRA under BioProject PRJNA1067836, BioSample SAMN45169522, processing folder is available upon request. Models and density maps for cryo-EM structures are deposited in the PDB (9ML4, 9ML5, 9ML6, and 9ML7) and maps are available on EMDB (48347, 48348, 48349, and 48350). Models and electron density maps for crystal structures were deposited in the PDB (9ML8 and 9ML9).

## Code availability

A Swift DMS program for processing and visualizing DMS data is available upon request.

## Acknowledgments

We thank Jost Vielmetter and the Caltech Beckman Institute Protein Expression Center for protein production, the Caltech Beckman Institute Beacon Center for Single Cell Biology for conducting Beacon experiments, Songye Chen and the Caltech Cryo-EM facility for cryo-EM data collection, and Jens Kaiser, staff at Stanford Synchrotron Radiation Lightsource, and the Caltech Molecular Observatory for X-ray data collection support. Cryo-Electron microscopy was performed in the Beckman Institute Resource Center for Transmission Electron Microscopy at Caltech. Use of the Stanford Synchrotron Radiation Lightsource, SLAC National Accelerator Laboratory, is supported by the U.S. Department of Energy, Office of Science, Office of Basic Energy Sciences under Contract No. DE-AC02-76SF00515. The SSRL Structural Molecular Biology Program is supported by the DOE Office of Biological and Environmental Research, and by the National Institutes of Health, National Institute of General Medical Sciences (P30GM133894). These studies were funded by the National Institutes of Health (NIH) P01-AI165075 (T.H., P.D.B., P.J.B.), the Caltech Merkin Institute (P.J.B.), Wellcome Leap (P.J.B.), the Coalition for Epidemic Preparedness Innovations (CEPI) (P.J.B.), and the Boehringer Ingelheim Fonds PhD fellowship (V.A.B.). P.D.B. is a Howard Hughes Medical Institute Investigator. This work was supported, in whole or in part, by the Bill & Melinda Gates Foundation grant INV-034638 (P.J.B.). Under the grant conditions of the Foundation, a Creative Commons Attribution 4.0 Generic License has already been assigned to the Author Accepted Manuscript version that might arise from this submission.

