## Supplementary material for "Cross-reactive sarbecovirus antibodies induced by mosaic RBD-nanoparticles": SI Appendix

**This PDF file includes:**

Supplemental Materials and Methods
Figures S1 to S6
Tables S1 to S3
SI References

**Supplemental Materials and Methods**

Mammalian cell lines

HEK293T (pseudovirus production) and HEK293T-hACE2 cells (pseudovirus neutralization assays) were cultured in Dulbecco's modified Eagle's medium (DMEM) supplemented with 10% heat-inactivated fetal bovine serum (FBS, Bio-Techne) and 1% Penicillin/Streptomycin (Gibco) and 1% L-Glutamine (Gibco). HEK-293T cells expressing high levels of hACE2 (kindly provided by Kenneth Matreyek, Case Western Reserve University), used for SHC014 pseudovirus neutralization assays, were cultured in DMEM supplemented with 10% heat-inactivated FBS (Bio-Techne), 1% Penicillin/Streptomycin (Gibco), 1% L-Glutamine (Gibco), 0.75 mg/mL puromycin dihydrochloride (Research Products International) and 2 mg/mL doxycycline hydrochloride (Thermo Fisher Scientific). HEK293T/ACE2cl.22 cells (1) (*in vitro* selection experiments) were cultured in DMEM supplemented with 10% FBS and 10 mg/mL gentamicin. Cell lines were cultured at 37 °C and 5% $CO_2$.

Expi293F cells for protein expression were maintained in Expi293 expression medium (Thermo Fisher Scientific). Transfections were performed using the Expi293 Expression System Kit and maintained under shaking at 130 rpm at 37 °C and 8% $CO_2$.

RBD expression and production of RBD-NPs

SpyTag003 (2)-tagged RBDs for conjugation and Avi-tagged RBDs for Beacon assays and ELISAs were expressed by transient transfection in Expi293F cells and purified as previously described (3, 4). Briefly, mammalian expression vectors encoding the RBDs of SARS-2 Beta (GenBank QUT64557.1), RaTG13-CoV (GenBank QHR63300), SHC014-CoV (GenBank KC881005), Rs4081-CoV (GenBank KY417143), pangolin17-CoV (GenBank QIA48632), RmYN02-CoV (GSAID EPI_ISL_412977), Rf1-CoV (GenBank DQ412042), W1V1-CoV (GenBank KF367457), SARS-2 Wuhan-Hu-1 (GenBank MN985325.1), SARS-2 Omicron BA.1 (GenBank UFO69278.1), SARS-2 Omicron BA.4/BA.5 (GenBank UPP14409.1), SARS-2 Omicron BA.5-S8 (5), SARS-2 Omicron XBB.1(EPI_ISL_15031899), SARS-2 Omicron XBB.1.5 (GenBank UZG29433.1), SARS-2 Omicron JN.1 (GenBank OR816965.1), SARS-2 Omicron KP.3 (EPI_SET_240529sa), RsSTT200-CoV (EPI_ISL_852605), SARS-1 (GenBank AAP13441.1), Yun11-CoV (GenBank JX993988), BtkY72-CoV (GenBank KY352407), Khosta-2-CoV (QVN46569.1), or BM-4831-CoV (GenBank NC014470) with the appropriate tag were used for transfection as described (3, 4, 6). Genes for RBDs used for Beacon assays were co-expressed with a plasmid encoding an ER-directed BirA enzyme for *in vivo* biotinylation (7) (kind gift of Michael Anaya, Caltech) using a 1:1 RBD to BirA plasmid ratio for transfections. Biotinylation was confirmed using a gel shift assay with streptavidin. RBDs were purified from transiently-transfected Expi293F supernatants by Ni-NTA and size-exclusion chromatography (SEC) and stored at 4˚C or -80˚C after flash freezing in liquid nitrogen (8).

A soluble SARS-2 Wuhan-Hu-1 spike trimer with 6P stabilizing mutations (8) was expressed and purified (9) for structural studies. Human IgG mAbs and human ACE-2 fused to human IgG Fc (hACE2-Fc) (10) were expressed and purified as described (9, 10). SpyCatcher003-mi3 NPs (11) were produced in *B. subtilis* (Ingenza, LTD) and purified as described (12).

RBD-NPs were generated by incubating SpyCatcher003-mi3 with a 2-fold molar excess (RBD to mi3 subunit) of SpyTagged RBD (either a single RBD for homotypic RBD-NPs or an equimolar mixture of eight RBDs for mosaic-8b RBD-NPs) overnight at room temperature in Tris-buffered saline (TBS). Conjugated RBD-NPs were separated from free RBDs by SEC as described (4). Concentrations of conjugated RBD-NPs are reported based on RBD content determined using a Bio-Rad Protein Assay.

mAbs for characterization assays

Rabbit mAbs that bind to SpyCatcher (anti-SC003 E1) and mi3 (anti-mi3 A3) were identified from memory B cells from rabbits immunized with mosaic-8b or homotypic RBD-NPs, respectively. Activated PBMCs were loaded on a Beacon chip and assayed for binding to SpyCatcher and mi3.

Candidate IgG-secreting B cells were exported and VH and VL gene segments were cloned into expression vectors encoding human constant domains.

Human mAbs that bind to Pang17, SHC014, Rs4081, Rf1, or RmYN02 RBD but not any of the other seven RBDs on mosaic-8b RBD-NPs (anti-Pang17, anti-SHC014, anti-Rs4081, Anti-Rf1 422, Anti-Rf1 425, Anti-Rf1 428, anti-RmYN02) were identified via contract by Biorad using their HuCAL technology with a positive selection and multiple negative selection strategy (Biorad). gBlocks corresponding to $V_H$ and $V_L$ sequences provided by Biorad were cloned into expression vectors encoding human constant domains.

A rabbit mAb (anti-RaTG13 F1) that binds to RaTG13 RBD and not the other seven RBDs on the mosaic-8b RBD-NPs was identified from a rabbit immunized with mosaic-8b RBD-NPs. Activated PBMCs were loaded on a Beacon chip and assayed for binding to the eight RBDs of mosaic-8b and the NP components (mi3 and SpyCatcher). Candidate IgG-secreting B cells were exported and VH and VL gene segments were cloned into expression vectors encoding human constant domains. Candidate mAbs were expressed and screened by ELISA for RaTG13 binding and minimal binding to other antigens.

β38 IgG (binds SARS-2 Beta RBD) and M8a-7 IgG (binds WIV1 RBD) were described previously (13, 14).

To confirm the appropriate specificity, anti-scaffold and anti-RBD mAbs were expressed and purified by MabSelect SURE affinity chromatography as described in the Ab cloning, protein expression, and purification section below. Purified mAbs were tested for binding to all eight RBDs (anti-RBD mAbs), or for binding to scaffold components (anti-SC003 and anti-mi3 mAbs) in ELISAs as described in the Binding and Competition ELISAs section below. Briefly, 2.5-5 mg of RBD, soluble SpyCatcher003, SC003-mi3, RBD-NPs, or mi3 was coated on MaxiSorp™ 384-well plates (Millipore Sigma) overnight. Excess RBD was removed by aspiration and plates were blocked with 3% bovine serum albumin (BSA). A dilution series of mAb was incubated for 3 hours, followed by incubation with an HRP-conjugated goat anti-human IgG (SouthernBiotech) secondary and detection by luminescence using SuperSignal ELISA Femto Maximum Sensitivity Substrate (Thermo Fisher Scientific). Experiments were conducted in duplicate (anti-RBD mAbs) or quadruplicate (anti-scaffold mAbs) with replicates averaged and standard deviations reported as error bars.

Anti-SC003 E1 and anti-mi3 A3 Fabs were generated from IgGs by incubating activated papain (Sigma-Aldrich) with 1-5 mg of IgG at a 1:100 ratio at 37 ºC for one hour. Fabs were separated from uncleaved IgG and papain by MabSelect SURE column and purified by SEC (Superdex 200 10/300; Cytiva).

A sandwich ELISA was used to characterize the NPs. Briefly, 2.5-5 mg of Fab was adsorbed to MaxiSorp™ 384-well plates (Millipore Sigma) overnight. Excess Fab was removed by aspiration and plates were blocked with 3% BSA. Plates were incubated with 0.25 mg/mL of mosaic-8b RBD-NP, homotypic RBD-NP, or SC003-mi3 for 2 hours at room temperature, followed by incubation with 1 µg/mL IgG for 2 hours. Bound IgG was detected with an Fc-specific secondary conjugated to HRP (goat anti-human IgG Fc; SouthernBiotech) and luminescence was read after incubation with SuperSignal ELISA Femto Maximum Sensitivity Substrate (Thermo Fisher Scientific). Experiments included four technical replicates, which were averaged for visualization.

Mosaic-8b RBD-NP rabbit immunizations

Immunizations were performed by Labcorp Drug Development using IACUC-approved protocols. All procedures in this study design are in compliance with the U.S. Department of Agriculture's (USDA) Animal Welfare Act (9 CFR Parts 1, 2, and 3); the Guide for the Care and Use of Laboratory Animals (Institute of Laboratory Animal Resources, National Academy Press, Washington, D.C., 2011); and the National Institutes of Health, Office of Laboratory Animal Welfare. Whenever possible, procedures in this study are designed to avoid or minimize discomfort, distress, and pain to animals. Five 7-8 week old New Zealand White female rabbits (Envigo) were immunized intramuscularly with 50 µg of conjugated mosaic-8b RBD-NPs (calculated as the mass of the RBD, assuming 100% efficiency of conjugation to SpyCatcher003-mi3, consistent with previous experimental results (3)) in 50% v/v AddaVax™ adjuvant (Invivogen). One female rabbit was immunized intramuscularly with 50 µg of homotypic SARS-2 RBD-NPs in 50% v/v AddaVax™ adjuvant. Animals received 250 µL per quadricep for a total 500 µL administration. Animals were boosted 4 weeks after the prime with a second dose of the same quantity of antigen in adjuvant. Peripheral blood mononuclear cells (PBMCs) and serum were obtained at day 0, 28, 42, 56, 90, and 119. PMBCs were stored at -80 ˚C until use.

Memory B cell activation

PMBCs from were thawed and MACS-enriched as described by Bruker Cellular Analysis in the rabbit Memory B cell workflow (Berkeley Lights, Incorporated). Briefly, PBMCs were thawed at 37 ˚C and drop-wise added to PBMC Thaw Media, centrifuged at 300 x *g* at 4°C, and then resuspended in MACS buffer supplemented with normal mouse serum (Thermo Fisher Scientific 10410) at 5% v/v for blocking. Memory B cells from PBMCs were enriched using MACS with biotinylated anti-IgG (anti-B cell receptor) Ab for labeling of the memory B cells and anti-biotin microbeads (Miltenyi 130-105-637) and MS columns (Miltenyi 130-042-201) for capture and recovery of the labeled cell population. Enriched memory B cells were activated in Activation Media (Bruker Cellular Analysis) for four before loading on the Beacon. PBMC thawing procedures, input cell density, enrichment conditions, and cell density for activation culture were followed from the Bruker Cellular Analysis workflow.

Bruker Cellular Analysis Beacon assays and cell export

Activated PBMCs were loaded on the Bruker Cellular Analysis Beacon instrument at ~8.0 x $10^6$ cells/mL. B cells were penned using opto-electero positioning (OEP) into individual nanopens on a 20k or an 11k OptoSelect Beacon microfluidic chip. Assays were conducted by providing a concentrated mixture of 5% goat anti-rabbit IgG (Fc) Coated Polystyrene Particles (Spherotech RPFc-60-5), 1:500 dilution of Alexa Fluor 488 goat anti-rabbit IgG (Jackson 111-546-144) and soluble biotinyated antigens coupled to streptavidin (SA)-fluorophore conjugates. Assay antigens were labeled with biotin via co-expression with BirA and purified and stored in aliquots as described above. Antigens were prepared for Beacon assays by labeling in 10 µg quantities with a 1:0.8 molar ratio of streptavidin-fluorphore conjugates. Conjugates were Streptavidin-Alexa 555 (PE, Thermo Fisher Scientific S32355), Streptavidin-Alexa 680 (Cy5, Thermo Fisher Scientific S32358), and Streptavidin Alexa 594 (TR, Thermo Fisher Scientific S32356) The labeling mixture was incubated at 4 ˚C overnight or until ready for use. The labeled stock was diluted to 20 µg/mL and then used in the assay bead mixture by supplementation to a final individual antigen concentration of 0.5 mg/mL.

Assay images were obtained every 7 minutes for 60 minutes. Multiple assays were run sequentially between 30-minute culture intervals using automated assay bead import. Nanopens were scored as positive or negative for IgG secretion and antigen capture using Bruker Cellular Analysis software for initial scoring and verification by manual inspection for each channel. Scored image sets were intersected to find cells of interest with binding observed for multiple antigens. Cells of interest were exported into 96-well plates containing RIPA (Thermo Fisher Scientific 89901) lysis buffer supplemented with RNAse inhibitor (New England Biolabs (NEB) M0134L) at a final concentration of 2U/µL and 5 mM DTT. This lysis solution was overlaid with mineral oil. Export plates were stored at -70 °C prior to cloning.

Ab cloning, protein expression, and purification

To recover Ig gene sequences of interest, RNA was purified from the single cell exports using RNAClean XP beads (Beckman Coulter A63987) and reverse transcribed into a cDNA library using reagents and guidelines provided by Bruker Cellular Analysis. Intermediate PCR products spanning $V_H$ and $V_L$ domains obtained from individual B cells were confirmed by agarose gel and sequenced by Plasmidsaurus. Gene blocks containing the coding sequences for rabbit $V_H$ and $V_L$ domains obtained from individual B cells were purchased from Integrated DNA Technologies (IDT).

Expression constructs were assembled using Golden Gate Assembly (NEB). The heavy chain for IgG expression was constructed by subcloning the $V_H$ gene into a p3BNC expression vector encoding human IgG1 $C_H1$, $C_H2$, and $C_H3$ domains. The expression plasmid for the light chain was constructed by subcloning the $V_L$ gene into a p3BNC vector that also encoded human kappa $C_L$.

IgGs were expressed in Expi293F cells by transient transfection and purified from cell supernatants using MabSelect SURE columns (Cytiva). Fabs were expressed as His-tagged truncated IgG heavy chains paired with the appropriate light chain and purified by Ni-NTA chromatography. IgGs and Fabs were further purified by SEC using a HiLoad 16/600 Superdex 200 column (Cytiva). Proteins were concentrated using a 30 kDa cutoff concentrator (EMD Millipore) to 10 to 15 mg/mL, and final concentrated proteins were stored at 4 ˚C.

Binding and Competition ELISAs

ELISAs were performed using a Tecan Evo liquid handling robot. For binding assays, Nunc® MaxiSorp™ 384-well plates (Millipore Sigma) were coated with 2.5-5.0 µg/mL of a purified RBD in 0.1 M $NaHCO_3$ pH 9.8 and stored overnight at 4 ˚C. Antigen was aspirated and plates were blocked with 3% bovine serum albumin (BSA) for an hour at room temperature. Blocking solution was removed and 4-fold serial dilutions of purified IgG or supernatant was added and incubated at room temperature for 3 hours. Plates were then washed with Tris-buffered saline with 0.1% Tween-20 (TBST) and incubated with secondary HRP-conjugated goat anti-human IgG (SouthernBiotech) at a 1:100,000 dilution for one hour at room temperature. Plates were washed with TBST, developed using SuperSignal ELISA Femto Maximum Sensitivity Substrate (Thermo Fisher Scientific), and luminescence was measured with a Tecan Infinite M1000 plate reader. ELISA data were collected in quadruplicates. Where indicated, curves were plotted and integrated to obtain half-maximal effective concentrations ($EC_{50}$) using Graphpad Prism v9.3.1 assuming a one-site binding model with a Hill coefficient. Data points represent the mean and error bars represent the standard deviation of the technical replicates.

Competition ELISAs were performed using a Tecan Evo liquid handling robot. Nunc® MaxiSorp™ 384-well plates (Millipore Sigma) were adsorbed with 5 µg/mL of Fab corresponding to mAb of known RBD epitope in 0.1 M $NaHCO_3$ pH 9.8 and stored overnight at 4 ˚C. Fab was removed and plates were blocked with 3% BSA in TBST for one hour at room temperature. Blocking solution was removed via aspiration and SARS-2 Wuhan-Hu-1 RBD (5 µg/mL) was added and incubated for 2 hours at room temperature. The RBD was removed via aspiration and 1 µg/mL IgG was added and incubated for 2 hours. The plate was washed with TBST and bound IgG was detected using horseradish peroxidase-conjugated Goat Anti-Human IgG Fc (SouthernBiotech) (1 hour, room temperature) and developed with SuperSignal ELISA Femto Substrate (Thermo Fisher Scientific). Luminescence was measured with a Tecan Infinite M1000 plate reader. Measurements were performed in technical quadruplicates and means are shown in a heat map.

Pseudovirus neutralization assays

SARS-2 Wuhan-Hu-1 D614G, SARS-2 VOCs, SARS-CoV, WIV1, SHC014, BtKY72/SARS-1 chimera (including mutations allowing human ACE2 binding (15)), and Khosta2/SARS-1 chimera pseudoviruses based on HIV lentiviral particles were prepared as described (16, 17). BtKY72/SARS-1 and Khosta2/SARS-1 chimeric spikes were constructed by replacing the RBD of the SARS-1 spike with the BtKY72 or Khosta2 RBD as described (4). Assays were done using 4-fold dilutions of purified IgGs at a starting concentration of 100 µg/mL by incubating with a pseudovirus at 37 ˚C for an hour. After incubating with $293T_{ACE2}$ target cells for 48 hours at 37 ˚C, cells were lysed with Luciferase Cell Culture Lysis 5x reagent (Promega). Luciferase activity in lysates was measured and relative luminescence units (RLUs) were normalized to values derived from cells infected with pseudovirus in the absence of IgG. Data were collected at each IgG concentration in duplicate and averaged. Half-maximal inhibitory concentrations ($IC_{50}$ values) were determined using nonlinear regression in AntibodyDatabase (18).

DMS

DMS studies used to map epitopes recognized by mAbs were performed in duplicates using SARS-2 Beta RBD libraries (generously provided by Tyler Starr, University of Utah) as described previously (19). RBD libraries were induced for RBD expression in galactose-containing synthetic defined medium with casamino acids (6.7g/L Yeast Nitrogen Base, 5.0 g/L Casamino acids, 1.065 g/L MES acid, 2% w/v galactose, and 0.1% w/v dextrose). After inducing for 18 hours, cells were washed 2x and then incubated with monoclonal Abs (dilutions chosen to give sub-saturating binding to RBDs) for 1 hour at RT with gentle agitation after which cells were washed 2x and labeled for 1 hour with secondary Ab: 1:200 Allophycocyanin-AffiniPure Goat Anti-Human IgG/Fcγ Fragment Specific (Jackson ImmunoResearch 109-135-098, RRID:AB_2337690) for human mAbs.

Stained yeast cells were processed on a Sony SH800 cell sorter and were gated to capture RBD mutants that had reduced mAb binding but relatively high RBD expression. For each sample, cells were collected up until around 3-5 x $10^6$ $RBD^+$ cells were processed (which corresponded to around 2 x $10^5$- 1 x $10^6$ $RBD^+$ Ab escaped cells). mAb-escaped cells were grown overnight in a synthetic defined medium with casamino acids, 100 U/mL penicillin, and 100 μg/mL streptomycin to expand cells prior to plasmid extraction. DNA extraction and Illumina sequencing were carried out as previously described (20). Raw sequencing data are available on the NCBI SRA under BioProject PRJNA1067836, BioSample SAMN45169522. Escape fractions were computed using processing steps described previously (20, 21) and implemented using a Swift DMS program (processing folder and program available from authors upon request). Escape scores were calculated with a filter to remove variants with deleterious mutations that escape binding due to poor expression, >1 amino acid mutation, or low sequencing counts as described (20, 22).

Logo plot visualizations of escape maps were created using Swift DMS (20), where letter height indicates the escape score for that amino acid mutation, and height of the stack of letters indicate the total site-wise escape metric, calculated as previously described (20). Letters for each site were colored according to epitope class. For structural visualizations, an RBD surface (PDB 6M0J) was colored by the site-wise escape metric at each site, with red scaled to be the maximum used to scale the y-axis for mAbs. Residues that exhibited the greatest escape fractions were highlighted with their residue number and colored according to epitope class.

*In vitro* selection experiments

To identify viral escape substitutions in the presence of mAb, we used a recombinant replication-competent vesicular stomatitis virus (rVSV) encoding the spike proteins of either SARS-2 Wuhan-Hu-1, SARS-2 VOC BA.2, SARS-2 VOC XBB.1.5, or SARS-1 as described (1, 23). Briefly, viral populations containing $10^6$ infectious units of either rVSV/SARS-2/$GFP_{2E1}$, rVSV/SARS-2/$GFP_{BA.2}$, rVSV/SARS-2/$GFP_{XBB.1.5}$, or rVSV/SARS-2/$GFP_{SARS-1}$ (1) were incubated with mAb at a concentration 10x above its $IC_{50}$ value for 1 hour at 37°C. Each virus/mAb mixture was then added to HEK-293T/ACE2cl.22 cells. In parallel, as a control, the four viral populations were added to the cells in the absence of any mAb. After 24 hours, the medium was replaced with fresh medium containing either a mAb or DMEM only. Following another 24 hours, the virus-containing supernatant was filtered through a 0.22 μm 96-well filter plate. The filtered supernatant (100 μL) was then incubated with the same concentration of rabbit mAb for 1 hour at 37°C, as described above. Each virus/mAb mixture was used to inoculate HEK-293T/ACE2cl.22 cells for a second passage (p2). Medium was again replenished with fresh mAb-containing medium or DMEM only after 24 hours, and the p2 viral populations were harvested after 48 hours. RNA was extracted from 100 μL of filtered p2 supernatant and reverse-transcribed using the SuperScript VILO cDNA Synthesis Kit (Thermo Fisher Scientific). Sequences encoding the extracellular domain of spike were amplified using KOD Xtreme Hot Start Polymerase (Sigma-Aldrich, 719753). Resulting PCR products were sequenced using Illumina MiSeq Nano 300 V2 cycle kits (Illumina, MS-103-1001) to identify mAb escape substitutions in the RBD. Specifically, sequencing reads were aligned to the corresponding RBD reference sequence and annotated for the presence of mutations. A variant was defined as occurring at a frequency >3% of reads at that position.

Cryo-EM Sample Preparation

Complexes of SARS-2 Wuhan-Hu-1 spike and Fabs were formed by incubating a purified spike trimer and a Fab at a molar ratio of 1:3.3 at room temperature for 30 minutes at a final spike concentration of ~2 mg/ml. Prior to freezing, fluorinated octylmaltoside solution (Anatrace) was added to the spike-Fab complex to a final concentration of 0.02% (w/v). Immediately after detergent addition, 3 µL of the spike-Fab complex/detergent mixture was applied to QuantiFoil 300 mesh 1.2/1.3 grids (Electron Microscopy Sciences) that had been freshly glow discharged for with 1 min at 20 mA using PELCO easiGLOW (Ted Pella). Grids were blotted with 0 blot force for 3 seconds with the Whatman No.1 filter paper at 100% humidity and room temperature before vitrification in 100% liquid ethane using a Mark IV Vitrobot (Thermo Fisher Scientific).

Cryo-EM data collection and processing
Single-particle cryo-EM datasets for complexes of SARS-2 Wuhan-Hu-1 spike 6P with M8b-A10 Fab, M8b-C9 Fab, and M8b-C10 Fab were collected using SerialEM (24) on a 300 keV Titan Krios (Thermo Fisher Scientific) equipped with a K3 direct electron detector camera (Gatan). Movies were recorded with a total dosage of 60 $e^-/Å^2$ and a defocus range of -1 to -3 µm in 40 frames using a 3x3 beam image shift pattern with 3 exposures per hole in super-resolution mode at a pixel size of 0.416 Å. A single-particle cryo-EM dataset for SARS-2 Wuhan-Hu-1 spike 6P–M8b-B8 Fab complex was collected with SerialEM (24) on a 200 keV Talos Arctica (Thermo Fisher Scientific) equipped with a K3 direct electron detector camera (Gatan). Movies were recorded with a total dosage of 60 $e^-/Å^2$ and a defocus range of -1 to -3 µm in 40 frames using a 3x3 beam image shift pattern with one single exposure per hole at a pixel size of 0.4345 Å. For all single-particle datasets, motion correction was performed with a binning factor of 2 using Patch Motion Correction in cryoSPARC v4.3 (25). Contrast Transfer Function (CTF) parameters were estimated via Patch CTF Estimation, and particles were picked with blob picker using a particle diameter of 100 to 200 Å in cryoSPARC v4.3 (25). Following inspection, particles were extracted and 2D classified in cryoSPARC v4.3 (25). After discarding bad particles, the remaining particles were used for *ab initio* modeling with 4 volumes, and these volumes were further refined with heterogeneous refinement in cryoSPARC v4.3 (25). Homogeneous and non-uniform refinements were carried out for the final reconstruction in cryoSPARC v4.3 (25). As the interactions between Fab and RBD were generally not well resolved when the RBDs adopted “up” conformations (26), we used local refinement to refine RBD-Fab regions as necessary. Masks used for local refinements were generated with UCSF Chimera (27) and local refinements were done in cryoSPARC v4.3 (25).

Cryo-EM Structure Modeling and Refinement
An initial model of the M8b-A10 Fab–SARS-2 RBD complex was generated by docking a mouse M8a-34 Fab–SARS-2 RBD (PDB 7UZD) into the locally refined cryo-EM density for M8b-A10 Fab–SARS-2 RBD using UCSF Chimera (27). The docked model was refined in Phenix (28) using real space refinement and the amino acid sequences for the Ab heavy and light chains were manually corrected in Coot (29). The M8b-A10 Fab–SARS-2 spike complex structure was generated by docking the refined structure of M8b-A10 Fab–SARS-2 RBD and a single-particle cryo-EM structure of SARS-2 spike trimer (PDB 7SC1) into the cryo-EM density map for the M8b-A10 Fab–SARS-2 spike complex in UCSF Chimera (27). The Fab-spike model was further refined in Phenix (28) using real space refinement, and outlier residues were rebuilt in Coot (29).The refined model of the M8b-A10 Fab was later used as the starting model for the remaining Fabs (M8b-B1, M8b-B8, M8b-C9 and M8b-C10) in the other Fab-spike trimer cryo-EM structures. Iterative real space refinements and model building were carried out separately in Phenix (28) and in Coot (29). Single-particle cryo-EM statistics are reported in *SI Appendix*, Table S2.

X-ray crystallography data collection, processing, and refinement
Crystallization trials for Fab–SARS-2 RBD complexes were set up using commercially available screens by mixing 0.2 µL of well solution and 0.2 µL of protein using sitting drop vapor diffusion (30) on a TTP LabTech Mosquito instrument at room temperature. Crystals for the M8b-B1–SARS-2 RBD were obtained from several conditions. A final dataset was collected from a crystal grown in a JCSG+ screen (Molecular Dimensions) containing 0.2 M lithium sulfate, 0.1 sodium citrate, and 20% (w/v) PEG 1,000. Crystals for the M8b-C9–SARS-2 RBD complex were also obtained from many conditions, and a final dataset was collected from a crystal grown in a PEGRx (Hampton.

Research), containing 2% v/v 1,4-dioxane, 0.1M Tris pH 8.0, and 15% (w/v) PEG 3,350. Crystals were cryoprotected in the crystallization solution mixed with 20% glycerol before flash-cooling in liquid nitrogen.

X-ray diffraction data were collected at 100 K at a wavelength of 0.9795 Å at the Stanford Synchrotron Radiation Lightsource (SSRL) beamline 12-2 using an EIGER 2XE 16M pixel array detector (Dectris) with the Blu-ice interface (31). X-ray datasets were indexed and integrated with XDS (32) and scaled with Aimless (33). The M8b-B1 Fab–SARS-2 RBD and M8b-C9 Fab–SARS-2 RBD structures were solved by molecular replacement using Phaser in Phenix (28). Iterative refinement and model-building cycles were carried out in Phenix (28) and Coot (29). The final refined M8b-B1–SARS-2 RBD structure included 97.8% of residues in Ramachandran favored regions and 2.2% in an allowed region. The final M8b-C9–SARS-2 RBD structure included 97.5% of residues in Ramachandran favored regions and 2.5% in an allowed region. Crystallographic statistics are reported in *SI Appendix*, Table S3.

Structural Motif Searches

To identify Ab/RBD structures with a YY motif in CDRH3 that interacts with RBD residues 378-382, we used the structure motif search service at RCSB (34) with PDB ID 7RKU; residues A51, A52, A53, A54, A55, G104, and G105; RMSD cutoff of 2 Å; and an Atom Pairing setting of All Atoms. A second structure motif search was done for a single CDRH3 tyrosine (corresponding to $Y98_{VH}$ in M8b-C9) and RBD residues 378-379 with PDB ID 7RKU; residues A51, A52, G104, and G105; with exchanges of G105 to all amino acids; RMSD cutoff of 1.5 Å; and an Atom Pairing setting of Backbone Atoms.

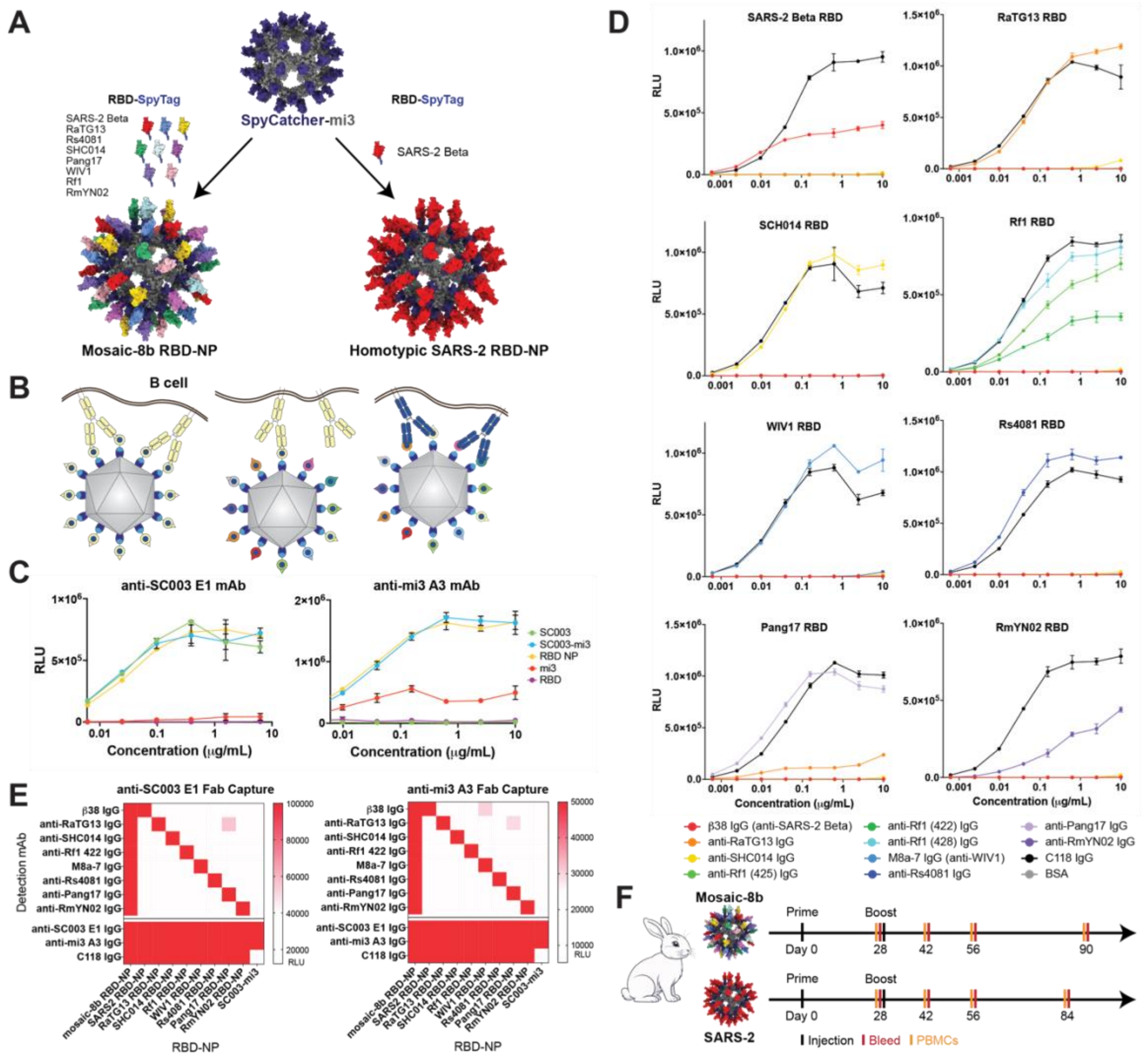

**Figure S1.** Experimental design for isolation of rabbit mAbs.

(A) Construction of mosaic-8b and homotypic SARS-2 RBD-NPs. Schematic of preparation of mosaic-8b and homotypic SARS-2 RBD-NPs. RBD-NP models were constructed with coordinates of an RBD (PDB 7BZ5), SpyCatcher (PDB 4MLI), and an i3-01 NP (PDB 7B3Y). (B) Hypothesis for preferential elicitation of cross-reactive Abs by mosaic-8b RBD-NPs versus homotypic SARS-2 RBD-NPs. Left: Both Fab arms of a membrane-bound B cell receptor recognizing a strain-specific variable epitope (yellow triangle) can bind to yellow antigens attached to a homotypic RBD-NP. Middle: Only one Fab arm of a B cell receptor recognizing a strain-specific variable epitope (yellow triangle) on a yellow antigen can bind to a mosaic NP when adjacent antigens are different. Right: Both Fab arms of a cross-reactive B cell receptor recognizing a conserved epitope (blue circles) can bind to adjacent antigens on a mosaic NP. (C) Anti-SC003 E1 mAb (left) and anti-mi3 A3 mAb (right) ELISAs of binding to components of NPs (soluble SpyCatcher003, unconjugated SpyCatcher-003 mi3 NPs, RBD-conjugated NPs, mi3, soluble RBD). Anti-SC003 E1 mAb recognizes SpyCatcher003 either as a soluble protein, attached to mi3 NPs, or on RBD-conjugated NPs. Anti-mi3 A3 mAb recognizes mi3 with and without SpyCatcher and RBD-conjugated NPs. Data are presented as the average of four replicates with error bars representing the standard deviation. (D) ELISAs of 8 anti-RBD mAbs show strain-specific binding to the RBDs present on mosaic-8b RBD-NPs. Data are presented as the average of two replicates with error bars representing the standard deviation. (E) Sandwich ELISA in which mosaic-8 RBD-NPs, each of the

8 homotypic RBD-NPs, and unconjugated SC003-mi3 NPs were captured on an ELISA plate by anti-SC003 E1 (left) or anti-mi3 A3 (right) and detected by strain-specific anti-RBD mAbs. High binding (red) indicates that the RBD corresponding to the mAb is present on the NPs. Low binding (white) indicates the RBD is absent. The human mAb C118 recognizes all 8 RBDs present on the mosaic-8b RBD-NPs (10). (F) Schematic of immunization regimen. Rabbits received RBD-NP prime and boost immunizations at days 0 and 28. Sera and PBMCs were collected at indicated days. Rabbit clipart from vecteezy.com.

**A**

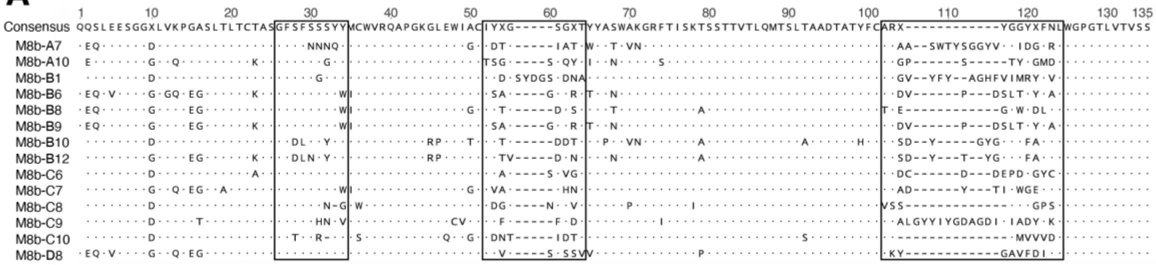

**B**

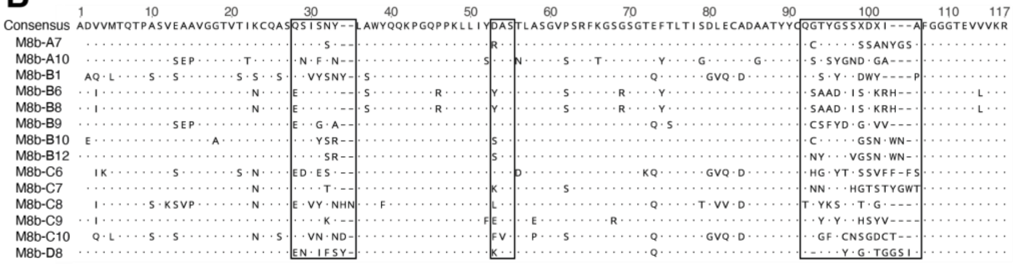

**C**

| mAb | Heavy Chain V-GENE | D-GENE | J-GENE | CDR1 Length | CDR2 Length | CDR3 Length | Light Chain V-GENE | J-GENE | CDR1 Length | CDR2 Length | CDR3 Length |
|---|---|---|---|---|---|---|---|---|---|---|---|
| **M8b-A7** | IGHV1S40 | IGHD6-1 | IGHJ4 | 9 | 8 | 21 | IGKV1S10 | IGKJ1-2 | 6 | 3 | 15 |
| **M8b-A10** | IGHV1S40 | IGHD7-1 | IGHJ6 | 9 | 9 | 12 | IGKV1S36 | IGKJ1-2 | 6 | 3 | 12 |
| **M8b-B1** | IGHV1S40 | IGHD6-1 | IGHJ4 | 9 | 9 | 19 | IGKV1S15 | IGKJ1-2 | 7 | 3 | 11 |
| **M8b-B6** | IGHV1S45 | IGHD1-1 | IGHJ4 | 9 | 9 | 14 | IGKV1S17 | IGKJ1-2 | 6 | 3 | 13 |
| **M8b-B8** | IGHV1S45 | IGHD4-2 | IGHJ4 | 9 | 8 | 11 | IGKV1S17 | IGKJ1-2 | 6 | 3 | 13 |
| **M8b-B9** | IGHV1S45 | IGHD1-1 | IGHJ4 | 9 | 9 | 14 | IGKV1S36 | IGKJ1-2 | 6 | 3 | 12 |
| **M8b-B10** | IGHV1S40 | IGHD6-1 | IGHJ4 | 9 | 8 | 16 | IGKV1S36 | IGKJ1-2 | 6 | 3 | 14 |
| **M8b-B12** | IGHV1S40 | IGHD6-1 | IGHJ4 | 9 | 8 | 16 | IGKV1S49 | IGKJ1-2 | 6 | 3 | 14 |
| **M8b-C6** | IGHV1S40 | IGHD2-1 | IGHJ2 | 9 | 9 | 14 | IGKV1S47 | IGKJ1-2 | 6 | 3 | 14 |
| **M8b-C7** | IGHV1S45 | IGHD4-1 | IGHJ4 | 9 | 8 | 14 | IGKV1S10 | IGKJ1-2 | 6 | 3 | 15 |
| **M8b-C8** | IGHV1S40 | IGHD1-1 | IGHJ4 | 8 | 9 | 10 | IGKV1S23 | IGKJ1-2 | 8 | 3 | 12 |
| **M8b-C9** | IGHV1S40 | IGHD6-1 | IGHJ4 | 9 | 8 | 23 | IGKV1S10 | IGKJ1-2 | 6 | 3 | 11 |
| **M8b-C10** | IGHV1S40 | IGHD7-1 | IGHJ4 | 8 | 8 | 9 | IGKV1S46 or IGKV1S60 | IGKJ1-2 | 7 | 3 | 13 |
| **M8b-D8** | IGHV1S45 | IGHD3-1 | IGHJ4 | 9 | 9 | 11 | IGKV1S10 | IGKJ1-2 | 6 | 3 | 14 |

**D**

| | A7 | A10 | B1 | B6 | B8 | B9 | B10 | B12 | C6 | C7 | C8 | C9 | C10 | D8 |
|---|---|---|---|---|---|---|---|---|---|---|---|---|---|---|
| **A7** | | 71.1 | 71.2 | 68.2 | 72.7 | 70.5 | 74 | 70.1 | 72.7 | 70.9 | 71.9 | 72.9 | 73.2 | 68.2 |
| **A10** | 71.1 | | 73.3 | 79.3 | 75 | 80.2 | 67.7 | 72.4 | 81.8 | 76 | 77.3 | 69.2 | 74.8 | 77.5 |
| **B1** | 71.2 | 73.3 | | 71 | 70.8 | 73.3 | 73.1 | 71 | 79.4 | 74 | 73.1 | 77.6 | 74.6 | 73.1 |
| **B6** | 68.2 | 79.3 | 71 | | 80.3 | 97.5 | 64.5 | 73.2 | 79.3 | 81 | 71.9 | 68.5 | 68.6 | 79.5 |
| **B8** | 72.7 | 75 | 70.8 | 80.3 | | 82.8 | 73 | 77.9 | 77.7 | 83.3 | 76.3 | 69.8 | 75.2 | 83.2 |
| **B9** | 70.5 | 80.2 | 73.3 | 97.5 | 82.8 | | 66.9 | 75.6 | 81.8 | 81.8 | 74.4 | 70.8 | 71.1 | 78.7 |
| **B10** | 74 | 67.7 | 73.1 | 64.5 | 73 | 66.9 | | 86.2 | 74.2 | 72.4 | 72.4 | 70.5 | 74.6 | 69.1 |
| **B12** | 70.1 | 72.4 | 71 | 73.2 | 77.9 | 75.6 | 86.2 | | 74.8 | 77 | 71.5 | 70.5 | 71.3 | 73.2 |
| **C6** | 72.7 | 81.8 | 79.4 | 79.3 | 77.7 | 81.8 | 74.2 | 74.8 | | 80.2 | 81 | 75.4 | 78.5 | 78.5 |
| **C7** | 70.9 | 76 | 74 | 81 | 83.3 | 81.8 | 72.4 | 77 | 80.2 | | 75.2 | 72.1 | 75.8 | 78.5 |
| **C8** | 71.9 | 77.3 | 73.1 | 71.9 | 76.3 | 74.4 | 72.4 | 71.5 | 81 | 75.2 | | 71.5 | 78.6 | 73.7 |
| **C9** | 72.9 | 69.2 | 77.6 | 68.5 | 69.8 | 70.8 | 70.5 | 70.5 | 75.4 | 72.1 | 71.5 | | 70.5 | 67.7 |
| **C10** | 73.2 | 74.8 | 74.6 | 68.6 | 75.2 | 71.1 | 74.6 | 71.3 | 78.5 | 75.8 | 78.6 | 70.5 | | 72 |
| **D8** | 68.2 | 77.5 | 73.1 | 79.5 | 83.2 | 78.7 | 69.1 | 73.2 | 78.5 | 78.5 | 73.7 | 67.7 | 72 | |

**E**

| | A7 | A10 | B1 | B6 | B8 | B9 | B10 | B12 | C6 | C7 | C8 | C9 | C10 | D8 |
|---|---|---|---|---|---|---|---|---|---|---|---|---|---|---|
| **A7** | | 75.4 | 73.9 | 80.7 | 80.7 | 81.6 | 90.4 | 90.4 | 77 | 88.6 | 71.3 | 86 | 72.8 | 85.2 |
| **A10** | 75.4 | | 67 | 74.1 | 74.1 | 80.2 | 74.3 | 75.2 | 67 | 73.7 | 69.6 | 73.9 | 67.9 | 74.8 |
| **B1** | 73.9 | 67 | | 68.1 | 68.1 | 74.1 | 74.6 | 73.7 | 77.9 | 71.3 | 75.9 | 75 | 81.3 | 72.2 |
| **B6** | 80.7 | 74.1 | 68.1 | | 100 | 77 | 77 | 79.8 | 71.4 | 80.7 | 70.8 | 78.8 | 70.5 | 76.7 |
| **B8** | 80.7 | 74.1 | 68.1 | 100 | | 77 | 77 | 79.8 | 71.4 | 80.7 | 70.8 | 78.8 | 70.5 | 76.7 |
| **B9** | 81.6 | 80.2 | 74.1 | 77 | 77 | | 81.4 | 81.6 | 75.9 | 78.1 | 76.8 | 80.4 | 72.3 | 82.8 |
| **B10** | 90.4 | 74.3 | 74.6 | 77 | 77 | 81.4 | | 94.7 | 74.3 | 84.2 | 72.8 | 83.2 | 71.7 | 84.3 |
| **B12** | 90.4 | 75.2 | 73.7 | 79.8 | 79.8 | 81.6 | 94.7 | | 74.3 | 87.7 | 72.8 | 86 | 72.6 | 84.5 |
| **C6** | 77 | 67 | 77.9 | 71.4 | 71.4 | 75.9 | 74.3 | 74.3 | | 73.5 | 71.1 | 75 | 73.5 | 74.6 |
| **C7** | 88.6 | 73.7 | 71.3 | 80.7 | 80.7 | 78.1 | 84.2 | 87.7 | 73.5 | | 69.6 | 82.5 | 72.8 | 82.6 |
| **C8** | 71.3 | 69.6 | 75.9 | 70.8 | 70.8 | 76.8 | 72.8 | 72.8 | 71.1 | 69.6 | | 72.3 | 74.3 | 72.2 |
| **C9** | 86 | 73.9 | 75 | 78.8 | 78.8 | 80.4 | 83.2 | 86 | 75 | 82.5 | 72.3 | | 70.5 | 79.3 |
| **C10** | 72.8 | 67.9 | 81.3 | 70.5 | 70.5 | 72.3 | 71.7 | 72.6 | 73.5 | 72.8 | 74.3 | 70.5 | | 71.1 |
| **D8** | 85.2 | 74.8 | 72.2 | 76.7 | 76.7 | 82.8 | 84.3 | 84.5 | 74.6 | 82.6 | 72.2 | 79.3 | 71.1 | |

**Figure S2.** Sequence alignment and identity of VH and VL gene segments from cloned rabbit mAbs.

(A and B) Amino acid sequence alignment of 14 rabbit (A) VH and (B) VL gene segments. CDRs (boxed) were identified using IMGT/HighV-Quest (35). (C) Gene segment usage and CDR lengths were determined for rabbit mAbs using IMGT/HighV-Quest (35). (D,E) Matrix of percent amino acid identity between rabbit (D) VH and (E) VL gene segments.

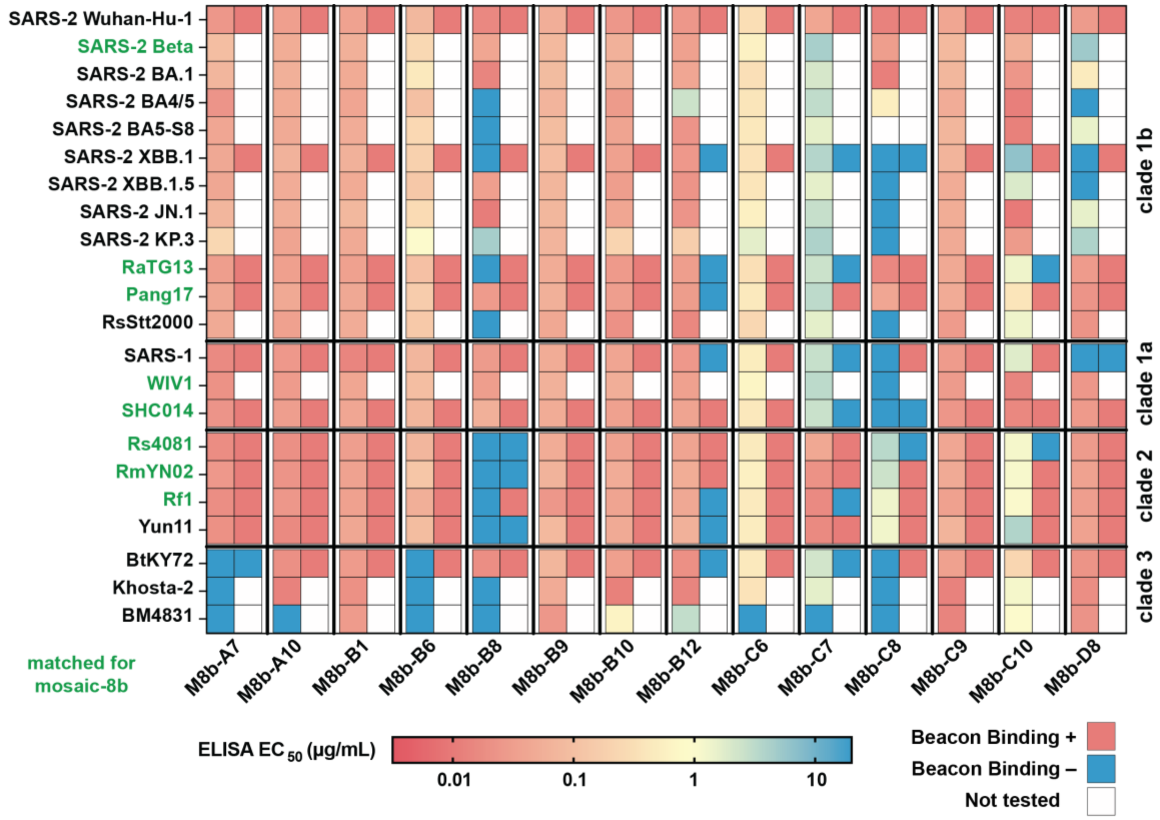

**Figure S3.** Comparison of ELISA data from recombinantly expressed mAbs and binding data from Beacon multiplexed assays.

For each mAb listed at the bottom, ELISA binding is depicted in the column on the left by a color gradient from blue (no binding) to coral (strong binding) for each RBD based on $EC_{50}$ values, and Beacon assay binding is depicted in the column on the right as binding (coral) or no binding (blue) based on detection of a bloom in the multiplexed assays for the indicated RBD. The Beacon multiplexed assays were validated by ELISAs of mAbs, with exceptions including nanopens that contained two cells rather than a single cell (M8b-C8), nanopens in which the signal could retrospectively be assigned to a cell in a neighboring nanopen (M8b-C7), or cells that appear to have died during the assays (M8b-B12).

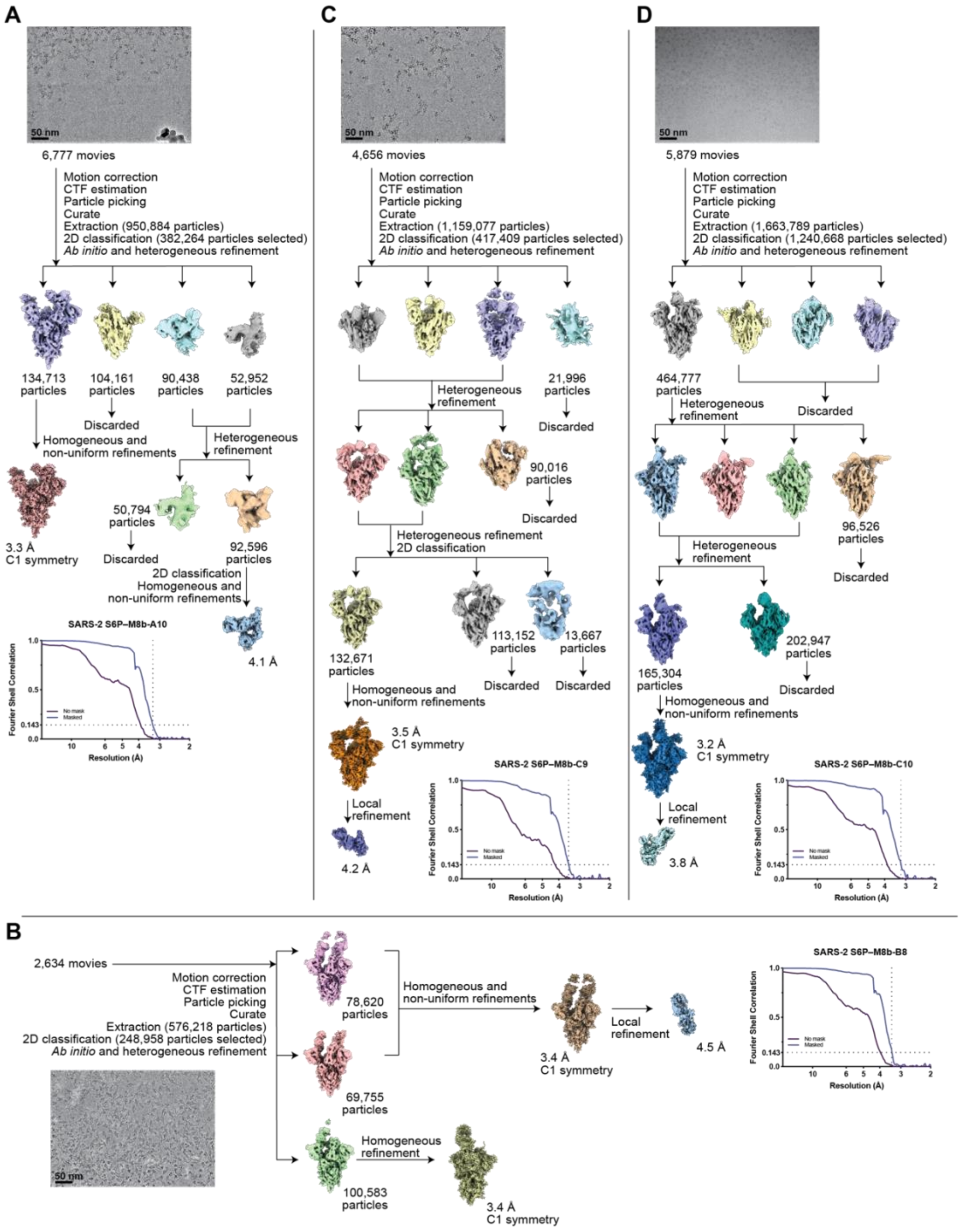

**Figure S4.** Single-particle cryo-EM processing for SARS-2 spike–mAb complexes.

Representative micrographs, workflows for single-particle data processing, and Fourier Shell Correlation plots for final reconstructions are shown for SARS-2 Wuhan-Hu-1 spike complexed with (A) M8b-A10, (B) M8b-B8, (C) M8b-C9, and (D) M8b-C10.

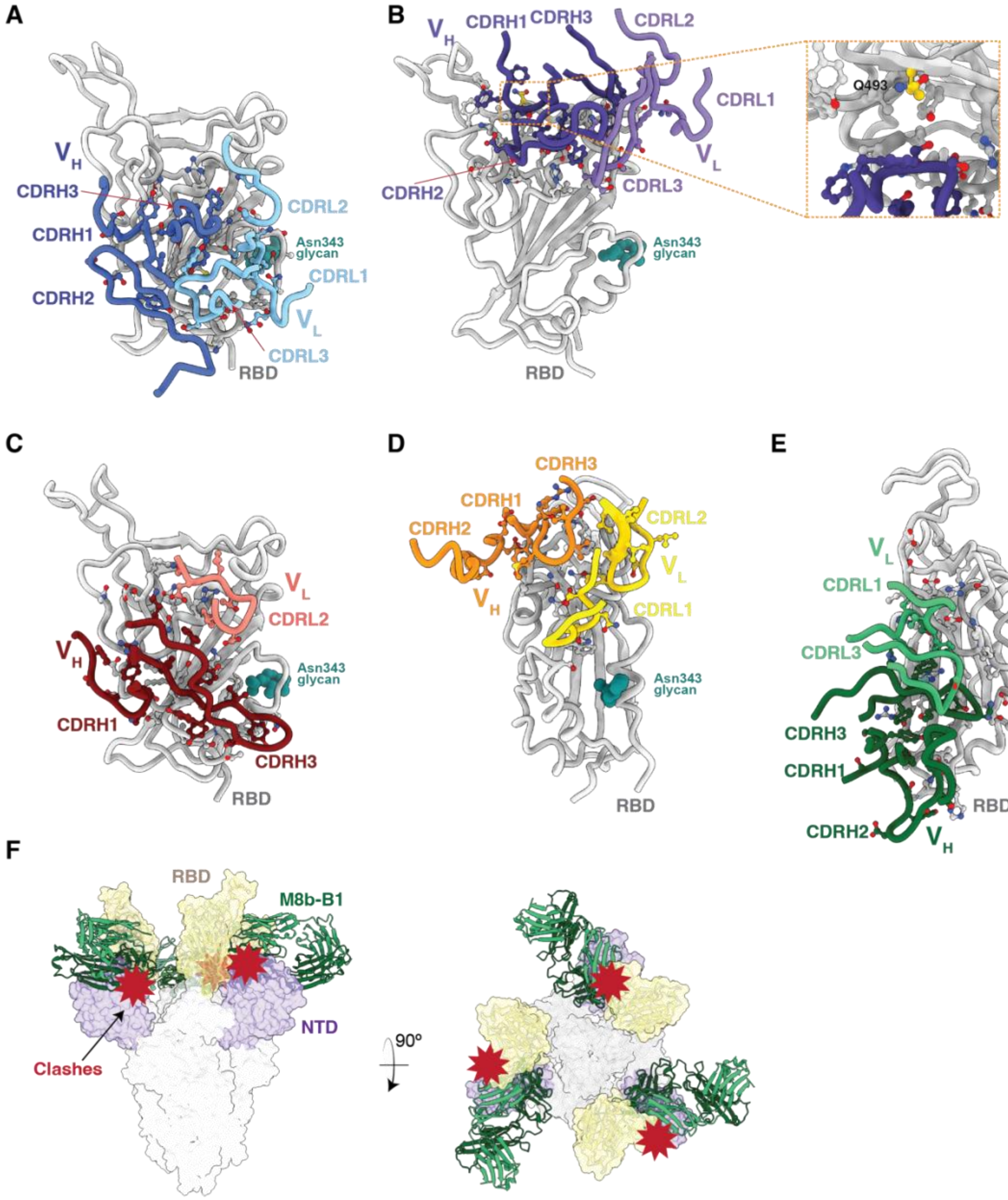

**Figure S5.** Rabbit mAb recognition of SARS-2 RBD.

Interactions are shown between the SARS-2 RBD and RBD-contacting CDRs of (A) M8b-A10, (B) M8b-B8, (C) M8b-C9, (D) M8b-C10, and (E) M8b-B1. RBD residue Q493 is highlighted in yellow in the panel b inset. (F) Modeling of the M8b-B1–RBD structure onto a SARS-2 spike trimer with all "up" RBDs (PDB 7SC1). SARS-2 spike is shown in surface representation with yellow RBDs and purple N-terminal domains (NTDs). The RBD portion of the M8bB1–RBD coordinates (cartoon representation; green M8b-B1 Fab, yellow RBD) was aligned on the RBDs of the spike trimer structure. Steric clashes between the M8b-B1 Fab and the spike NTD are highlighted as red bursts.

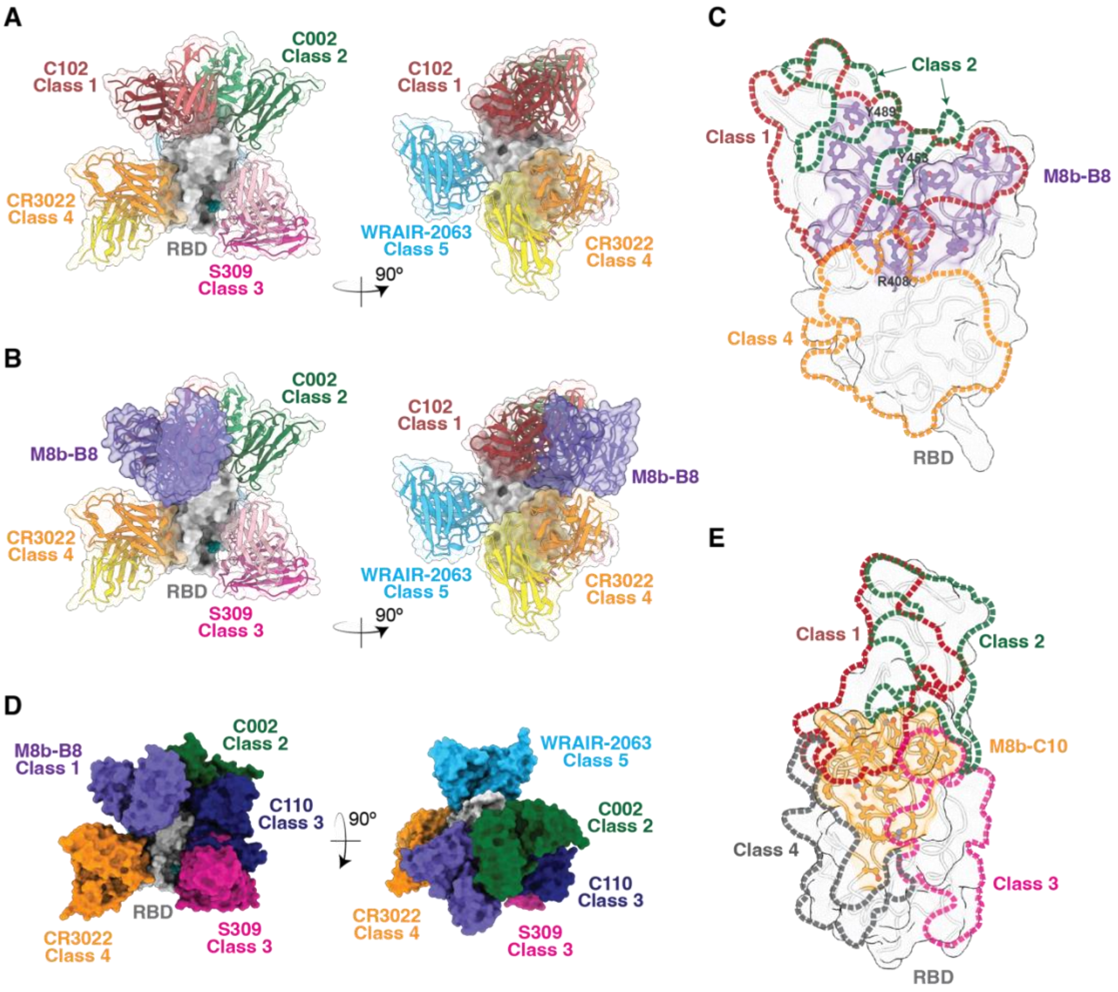

**Figure S6.** Epitope classifications of M8b-B8 and M8b-C10.

(A) $V_H$-$V_L$ domains of representative mAbs recognizing previously-defined RBD epitopes (9, 36, 37): C102: PDB 7K8M; C002: PDB 7K8T, S309: PDB 7JX3; CR3022: PDB 7LOP; WRAIR-2063: PDB 8EOO. (B) Overlap of the binding footprints of M8b-B8 (purple RBD residues), C102 (class 1, red dotted outline), C002 (class 2, green dotted outline), and CR3022 (class 4, orange dotted outline). (C) The M8b-B8 epitope (light purple) primarily overlaps with the class 1 RBD epitope (exemplified by C102; red dotted line). Two residues within the M8b-B8 epitope (Y453 and Y489) overlap with the class 2 RBD epitope (exemplified by C002; green dotted line), and one residue (Y408) overlaps with the class 4 RBD epitope (exemplified by CR3022; orange dotted line). (D) Composite model showing that binding of M8b-B8 could sterically interfere with the binding of the class 2 mAb C002, the class 3 mAb C110, and the class 4 mAb CR3022. (E) The epitope of M8b-C10 (orange) primarily overlaps with the class 1 RBD epitope (exemplified by C102; red dotted lines) and the class 3 RBD epitope (exemplified by S309; magenta dotted line).

**Table S1.** $V_H$ and $V_L$ sequences of anti-NP and strain-specific anti-RBD mAbs.

| mAb Name | VH/VL Origin | VH nucleotide sequence | VH amino acid sequence | VL nucleotide sequence | VL amino acid sequence | Reference |
|---|---|---|---|---|---|---|
| anti-SC003 E1 IgG | Rabbit | CAGCAGCTGGAGCAGTCCGAGGGAGACCTGGTCAAGCCTGGGGCATCCCTGACACTCTCCTGCAAAGCCTCTGGATTCTCCTTCAGTGGCAATGTTATGTGCTGGGTCCGCCAGGCTCCAGGGAAGGGGCTGGAGTGGATCGGATGCATTTATATTGTTGATGGCAAGACTTACTACGCGAGCTGGGTGAATGGCCGATTCACCATCTCCAAAACCTCGTCGACCACGGTGACTCTGCAAATGACCAGTCTGACAGCCGCGGACACGGCCACCTATTTCTGTGCGAGGGGGACTTATGCTGGTTATGGTGGTGGTGGTGATCTCTTTGACGAGTTTAATTTGTGGGGCCCCGGCACCCCGGTCACCGTCTCCTCA | QQLEQSEGDLVKPGASLTLSCKASGFSFSGNVMCWVRQAPGKGLEWIGCIYIVDGKTYYASWVNGRFTISKTSSTTVTLQMTSLTAADTATYFCARGTYAGYGGGDLFDEFNLWGPGTPVTVSS | GCCCTTGTGATGACCCAGACTCCAGCCTCCGTGTCTGAACCTGTGGGAGGCACAGTCACCGTCAAGTGCCAGGCCAGTGAGAACATTTATAGCAATTTAGCCTGGTATCAGCAGAAACCAGGGCAGCCTCCCAAGCTCCTGATCTACAAGGCATCCACTCTGGCATCTGGGGTCCCATCGCGGTTCAAAGGCAGTGGATCTCGGACAGAGTACACTCTCACCATCAGCGACCTGGAGTGTGCCGATGCTGCCACTTACTACTGTCAAAGCAATTTTGGTGGTACTAGTATGAGTTATTATTTCGGCGGATGGACCGAGGTGGTGGTCAAACGT | ALVMTQTPASVSEPVGGTVTVKCQASENIYSNLAWYQQKPGQPPKLLIYKASTLASGVPSRFKGSGSRTEYTLTISDLECADAATYYCQSNFGGTSMSYYFGGWTEVVVKR | This paper |
| anti-mi3 A3 IgG | Human | CAGCAGCTGGAGCAGTCCGGGGGAGGCCTGGTCCAGCCTGAGGGATCCCTGACACTCACCTGCAAAGCCTCTGGAATCGACTTCAGTAGCTACTACTACATGTGCTGGGTCCGCCAGGCTCCAGGGAAGGGGCTGGAGTTGATCGGATGCATTAATACTGGTGGTAATGGTGCCACTTACTACGCGAGCTGGGCGAAAGGCCGATTCACCATCTCCAGAACCTCGTCGACCACGGTGACTCTGCAAATGACCAGTTTGACAGCCGCGGACACGGCCACCTATTTCTGTGCGAGAGATCTCTATTATAATGTTGGTAACTATGGGGGCTTGTGGGGCCCAGGCACCCTGGTCACCGTCTCCTCA | QQLEQSGGGLVQPEGSLTLTCKASGIDFSSYYYMCWVRQAPGKGLELIGCINTGGNGATYYASWAKGRFTISRTSSTTVTLQMTSLTAADTATYFCARDLYYNVGNYGGLWGPGTLVTVSS | GCCCTTGTGATGACCCAGACTCCAGCCTCCGTGTCTGCCGCTGTGGGAGGCACAGTCACCATCAAGTGCCAGGCCAGTCAGAACATTAAGAACCAACTATCCTGGTATCAACAGAAACCAGGGCAGCGTCCCAAGCTCCTGGTCTACGATGCATCCGAACTGGCATCTGGGGTCCCATCACGGTTCGGTGGCAGTGGATCTGGGACAGAGTACACTCTCGCCATCAGCGACCTGGAGTGTGCCGATGCTGCCACTTACTACTGTCAACAGGGTTGGAATATTGATGGGGTTGATAATGTTTTCGGCGGAGGGACCGAGGTGGTGGTCAAA | ALVMTQTPASVSAAVGGTVTIKCQASQNIKNQLSWYQQKPGQRPKLLVYDASELASGVPSRFGGSGSGTEYTLAISDLECADAATYYCQQGWNIDGVDNVFGGGTEVVVK | This paper |
| Beta-38 IgG | Human | GAAGTGCAGCTGGTTCAGTCTGGCGCCGAAGTGAAGAAGCCTGGCGAGAGCCTGAAGATCAGCTGCAAAGGCAGCGGCTACAGCTTCACCAACTACTGGATCGGCTGGGTCCGACAGATGCCTGGCAAAGGCCTTGAGTGGATGGGCATCATCTACCCCGGCGATAGCGGCACCAGATACAGCCCTAGCTTTCAGGGCCAAGTGACCATCAGCGCCGACAAGAGCATCAGAACCGCCTACCTGCAGTGGTCCAGCCTGAAGGCCTCTGACAGCGCCATGTACTACTGCGCCAGATCTAGAGTGGGAGCCACCGGCGGCTACTACGACTACTACATGGATGTGTGGGGCCAGGGCACCACCGTGACAGTTTCTTCT | EVQLVQSGAEVKKPGESLKISCKGSGYSFTNYWIGWVRQMPGKGLEWMGIIYPGDSGTRYSPSFQGQVTISADKSIRTAYLQWSSLKASDSAMYYCARSRVGATGGYYDYYMDVWGQGTTVTVSS | CAGTCTGTGCTGACACAGCCTCCATCTGCCTCTGGCACACCTGGCCAGAGAGTGACCATCAGCTGTAGCGGCAGCAGCAGCAATCTCGGCGGCAACACCGTGAACTGGTATCAGCAGCTGCCTGGCACAGCCCCTAAACTGCTGATCTACAGCAACAACCAGCGGCCTAGCGGCGTGCCCGATAGATTTTCTGGCAGCAAGAGCGGCACAAGCGCCAGCCTGGCTATTTCTGGACTGCAGAGCGAGGACGAGGCCGACTATTATTGTGCCGCCTGGGACGACAGCCTGAACGGCCCTGTTTTTGGCACCGGCACCAAAGTGACAGTGCTG | QSVLTQPPSASGTPGQRVTISCSGSSSNLGGNTVNWYQQLPGTAPKLLIYSNNQRPSGVPDRFSGSKSGTSASLAISGLQSEDEADYYCAAWDDSLNGPVFGTGTKVTVL | Liu et al., Cell Host & Microbe, 2022 |
| anti-Pang17 IgG | Human | CAGGTGCAATTGGTGCAGAGCGGTGCCGAAGTGAAAAAACCGGGCAGCAGCGTGAAAGTTAGCTGCAAAGCATCCGGAGGGACGTTTAACGACTACGCTATGTCTTGGGTGCGCCAGGCCCCTGGACAGGGCCTCGAGTGGATGGGCGGTATCATCCCGAAATTCGGCACTGCGTACTACGCCCAGAAATTTCAGGGCCGGGTGACCATTACCGCCGATGAAAGCACCAGCACCGCCTATATGGAACTGAGCAGCCTGCGCAGCGAAGATACGGCCGTGTATTATTGCGCGCGTGGTCCGGCTGTTGGTTACCCGTCTAACATCTACTACGAATACCCGTACGATGTTTGGGGCCAAGGCACCCTGGTGACTGTTAGCTCA | QVQLVQSGAEVKKPGSSVKVSCKASGGTFNDYAMSWVRQAPGQGLEWMGGIIPKFGTAYYAQKFQGRVTITADESTSTAYMELSSLRSEDTAVYYCARGPAVGYPSNIYYEYPYDVWGQGTLVTVSS | GATATCCAGATGACCCAGAGCCCGAGCAGCCTGAGCGCCAGCGTGGGCGATCGCGTGACCATTACCTGCAGAGCCAGCCAGGACATTTCTAACCGTCTGGCTTGGTACCAGCAGAAACCGGGCAAAGCGCCGAAACTATTAATCTACGCTGCTTCTAACCTGCAAAGCGGCGTGCCGAGCCGCTTTAGCGGCAGCGGATCCGGCACCGATTTCACCCTGACCATTAGCTCTCTGCAACCGGAAGACTTTGCGACCTATTATTGCCATCAGTACTACTCTACTCCGCCGACCTTTGGCCAGGGCACGAAAGTTGAAATTAAA | DIQMTQSPSSLSASVGDRVTITCRASQDISNRLAWYQQKPGKAPKLLIYAASNLQSGVPSRFSGSGSGTDFTLTISSLQPEDFATYYCHQYYSTPPTFGQGTKVEIK | This paper |
| anti-Rf1 422 IgG | Human | GAAGTGCAATTGCTGGAAAGCGGCGGTGGCCTGGTGCAGCCGGGTGGCAGCCTGCGTCTGAGCTGCGCGGCGTCCGGATTCACCTTTTCTTCTGCTGCTATGCATTGGGTGCGCCAGGCCCCGGGCAAAGGTCTCGAGTGGGTTTCCTCTATCTCTTACTACGGTTCTTCTACCGACTATGCGGATAGCGTGAAAGGCCGCTTTACCATCAGCCGCGATAATTCGAAAAACACCCTGTATCTGCAAATGAACAGCCTGCGTGCGGAAGATACGGCCGTGTATTATTGCGCGCGTAACTTCCCGGGTTACGGTGCTTACTACATGGATTACTGGGGCCAAGGCACCCTGGTGACTGTTAGCTCA | EVQLLESGGGLVQPGGSLRLSCAASGFTFSSAAMHWVRQAPGKGLEWVSSISYYGSSTDYADSVKGRFTISRDNSKNTLYLQMNSLRAEDTAVYYCARNFPGYGAYYMDYWGQGTLVTVSS | GATATCCAGATGACCCAGAGCCCGAGCAGCCTGAGCGCCAGCGTGGGCGATCGCGTGACCATTACCTGCAGAGCCAGCCAGTCTATTGGTGACTACCTGAACTGGTACCAGCAGAAACCGGGCAAAGCGCCGAAACTATTAATCTACGCTGCTTCTTCTCTGCAAAGCGGCGTGCCGAGCCGCTTTAGCGGCAGCGGATCCGGCACCGATTTCACCCTGACCATTAGCTCTCTGCAACCGGAAGACTTTGCGACCTATTATTGCCAGCAGGCTTACTTCACTCCGCTGACCTTTGGCCAGGGCACGAAAGTTGAAATTAAA | DIQMTQSPSSLSASVGDRVTITCRASQSIGDYLNWYQQKPGKAPKLLIYAASSLQSGVPSRFSGSGSGTDFTLTISSLQPEDFATYYCQQAYFTPLTFGQGTKVEIK | This paper |
| anti-Rf1 425 IgG | Human | CAGGTGCAATTGGTGCAGAGCGGTGCCGAAGTGAAAAAACCGGGCAGCAGCGTGAAAGTTAGCTGCAAAGCATCCGGAGGGACGTTTTCTGACTACGCTATCTCTTGGGTGCGCCAGGCCCCTGGACAGGGCCTCGAGTGGATGGGCCGTATCTACCCGATCTTCGGCACTGCGAACTACGCCCAGAAATTTCAGGGCCGGGTGACCATTACCGCCGATGAAAGCACCAGCACCGCCTATATGGAACTGAGCAGCCTGCGCAGCGAAGATACGGCCGTGTATTATTGCGCGCGTGAACAGTACTACTCTTCTTCTATCGGTTTCGATTACTGGGGCCAAGGCACCCTGGTGACTGTTAGCTCA | QVQLVQSGAEVKKPGSSVKVSCKASGGTFSDYAISWVRQAPGQGLEWMGRIYPIFGTANYAQKFQGRVTITADESTSTAYMELSSLRSEDTAVYYCAREQYYSSSIGFDYWGQGTLVTVSS | GATATCGAACTGACCCAGCCGCCGAGCGTGAGCGTGAGCCCGGGCCAGACAGCTAGCATTACCTGTAGCGGCGATAAAATCCCGAAAAAAGCTGTTCATTGGTACCAGCAGAAACCGGGCCAGGCGCCGGTGCTGGTGATCTACGACGACAACGACCGTCCGAGCGGCATCCCGGAACGTTTTAGCGGATCCAACAGCGGCAACACCGCGACCCTGACCATTAGCGGCACCCAGGCGGAAGACGAAGCGGATTATTACTGCGCTTCTTGGGACCTGGGTTCTTACGTGTTTGGCGGCGGCACGAAGTTAACCGTTCTT | DIELTQPPSVSVSPGQTASITCSGDKIPKKAVHWYQQKPGQAPVLVIYDDNDRPSGIPERFSGSNSGNTATLTISGTQAEDEADYYCASWDLGSYVFGGGTKLTVL | This paper |
| anti-Rf1 428 IgG | Human | CAGGTGCAATTGGTGCAGAGCGGTGCCGAAGTGAAAAAACCGGGCAGCAGCGTGAAAGTTAGCTGCAAAGCATCCGGAGGGACGTTTTCTTCTTACGCTATCTCTTGGGTGCGCCAGGCCCCTGGACAGGGCCTCGAGTGGATGGGCGGTATCGTTCCGATCTTCGGCGCTGCGACTTACGCCCAGAAATTTCAGGGCCGGGTGACCATTACCGCCGATGAAAGCACCAGCACCGCCTATATGGAACTGAGCAGCCTGCGCAGCGAAGATACGGCCGTGTATTATTGCGCGCGTGACCTGCATTACTACGCTATCGATGTTTGGGGCCAAGGCACCCTGGTGACTGTTAGCTCA | QVQLVQSGAEVKKPGSSVKVSCKASGGTFSSYAISWVRQAPGQGLEWMGGIVPIFGAATYAQKFQGRVTITADESTSTAYMELSSLRSEDTAVYYCARDLHYYAIDVWGQGTLVTVSS | GATATCCAGATGACCCAGAGCCCGAGCAGCCTGAGCGCCAGCGTGGGCGATCGCGTGACCATTACCTGCAGAGCCAGCCAGGACATTATCACTTACCTGGCTTGGTACCAGCAGAAACCGGGCAAAGCGCCGAAACTATTAATCTACGCTGCTTCTACTCTGCAAAGCGGCGTGCCGAGCCGCTTTAGCGGCAGCGGATCCGGCACCGATTTCACCCTGACCATTAGCTCTCTGCAACCGGAAGACTTTGCGACCTATTATTGCCAGCAGTACGACCAGGGTCCGATCACCTTTGGCCAGGGCACGAAAGTTGAAATTAAA | DIQMTQSPSSLSASVGDRVTITCRASQDIITYLAWYQQKPGKAPKLLIYAASTLQSGVPSRFSGSGSGTDFTLTISSLQPEDFATYYCQQYDQGPITFGQGTKVEIK | This paper |
| anti-RmYN02 IgG | Human | CAGGTGCAATTGGTGCAGAGCGGTGCCGAAGTGAAAAAACCGGGCAGCAGCGTGAAAGTTAGCTGCAAAGCATCCGGAGGGACGTTTAACTCTTACTCTATCTCTTGGGTGCGCCAGGCCCCTGGACAGGGCCTCGAGTGGATGGGCGGTATCATCCCGATCTTCGGCACTGCGAACTACGCCCAGAAATTTCAGGGCCGGGTGACCATTACCGCCGATGAAAGCACCAGCACCGCCTATATGGAACTGAGCAGCCTGCGCAGCGAAGATACGGCCGTGTATTATTGCGCGCGTGACCTGGTTGACAACGCTGTTTCTATGTACCCGCTGGATCCGTGGGGCCAAGGCACCCTGGTGACTGTTAGCTCA | QVQLVQSGAEVKKPGSSVKVSCKASGGTFNSYSISWVRQAPGQGLEWMGGIIPIFGTANYAQKFQGRVTITADESTSTAYMELSSLRSEDTAVYYCARDLVDNAVSMYPLDPWGQGTLVTVSS | GATATCCAGATGACCCAGAGCCCGAGCAGCCTGAGCGCCAGCGTGGGCGATCGCGTGACCATTACCTGCAGAGCCAGCCAGGACATTCGTAACTACCTGGCTTGGTACCAGCAGAAACCGGGCAAAGCGCCGAAACTATTAATCTACGACGCTTCTAACCTGCAAAGCGGCGTGCCGAGCCGCTTTAGCGGCAGCGGATCCGGCACCGATTTCACCCTGACCATTAGCTCTCTGCAACCGGAAGACTTTGCGACCTATTATTGCCAGCAGTACGGTGCTACTTCTTACACCTTTGGCCAGGGCACGAAAGTTGAAATCAAA | DIQMTQSPSSLSASVGDRVTITCRASQDIRNYLAWYQQKPGKAPKLLIYDASNLQSGVPSRFSGSGSGTDFTLTISSLQPEDFATYYCQQYGATSYTFGQGTKVEIK | This paper |
| anti-Rs4081 IgG | Human | GAAGTGCAATTGGTGGAAAGCGGCGGTGGCCTGGTGAAACCAGGCGGCAGCCTGCGCCTGAGCTGCGCCGCCTCCGGATTCACCTTTAACTCTTACGCTCTGCATTGGGTGCGCCAGGCCCCGGGCAAAGGTCTCGAGTGGGTGGGCCGTATCCGTTCTAAAACTGACGGTGGTACTACTCATTATGCCGCCCCAGTGAAAGGCCGCTTTACCATTAGCCGCGATGATTCGAAAAACACCCTGTATCTGCAAATGAACAGCCTGAAAACCGAAGATACGGCCGTGTATTATTGCGCGCGTGGTGGTCTGTACGACTACTACTACGACTACTTCGATTACTGGGGCCAAGGCACCCTGGTGACTGTTAGCTCA | EVQLVESGGGLVKPGGSLRLSCAASGFTFNSYALHWVRQAPGKGLEWVGRIRSKTDGGTTHYAAPVKGRFTISRDDSKNTLYLQMNSLKTEDTAVYYCARGGLYDYYYDYFDYWGQGTLVTVSS | GATATCGTGCTGACCCAGCCGCCGAGCGTGAGCGGTGCACCGGGACAGCGAGTGACCATTAGCTGTAGCGGCAGCAGCAGCAACATTGGTATCAACACTGTGAACTGGTACCAGCAGCTGCCGGGCACGGCGCCGAAAGTTCTGATCTACTCTAACTCTCATCGCCCGAGCGGCGTGCCGGATCGCTTTAGCGGATCCAAAAGCGGCACCAGCGCCAGCCTGGCGATTACCGGCCTGCAAGCAGAAGACGAAGCGGATTATTACTGCTCTTCTACTGCTGAACTGGACTCTGGTTGGGTGTTTGGCGGCGGCACGAAGTTAACCGTTCTT | DIVLTQPPSVSGAPGQRVTISCSGSSSNIGINTVNWYQQLPGTAPKVLIYSNSHRPSGVPDRFSGSKSGTSASLAITGLQAEDEADYYCSSTAELDSGWVFGGGTKLTVL | This paper |
| anti-SCH014 IgG | Human | CAGGTGCAATTGGTGCAGAGCGGTGCCGAAGTGAAAAAACCGGGCAGCAGCGTGAAAGTTAGCTGCAAAGCATCCGGAGGGACGTTTTCTTCTTACGCTATCTCTTGGGTGCGCCAGGCCCCTGGACAGGGCCTCGAGTGGATGGGCGGTATCATCCCGGTTTTCGGCGAAGCGAACTACGCCCAGAAATTTCAGGGCCGGGTGACCATTACCGCCGATGAAAGCACCAGCACCGCCTATATGGAACTGAGCAGCCTGCGCAGCGAAGATACGGCCGTGTATTATTGCGCGCGTATGGGTTGGTGGGACTTCTGGGGCCAAGGCACCCTGGTGACTGTTAGCTCA | QVQLVQSGAEVKKPGSSVKVSCKASGGTFSSYAISWVRQAPGQGLEWMGGIIPVFGEANYAQKFQGRVTITADESTSTAYMELSSLRSEDTAVYYCARMGWWDFWGQGTLVTVSS | GATATCGAACTGACCCAGCCACCGAGCGTGAGCGTGAGCCCGGGCCAGACCGCGAGCATTACCTGTAGCGGCGATAACATCGGTCAGAAATCTTACAACTGGTACCAGCAGAAACCGGGCCAGGCGCCGGTGCTGGTGATCTACGACAACGACGACCGTCCGAGCGGCATCCCGGAACGTTTTAGCGGATCCAACAGCGGCAACACCGCGACCCTGACCATTAGCGGCACCCAGGCGGAAGACGAAGCGGATTATTACTGCCAGTCTTGGACTTGGTCTTGGCTGTCTTACGTGTTTGGCGGCGGCACGAAGTTAACCGTTCTT | DIELTQPPSVSVSPGQTASITCSGDNIGQKSYNWYQQKPGQAPVLVIYDNDDRPSGIPERFSGSNSGNTATLTISGTQAEDEADYYCQSWTWSWLSYVFGGGTKLTVL | This paper |
| anti-RaTG13 F1 | Rabbit | CAGCAGCTGGAGCAGTCCGGGGGAGGCCTGGTCCAGCCTGAGGGATCCCTGACACTCACCTGCACAGTCTCTGGATTCTCGTGGAAGTACGATTACATGTGCTGGGTCCGCCAGGCTCCAGGGAAGGGGCTGGAGTGGATCGTATGTATTTATGGTGATAGTGGTAGCACATGGTACGCGAGCTGGGTGAATGGCCGATTCACCATTTCCAAAACCTCGTCGACCACGGTGACTCTGCAAATGGCTAGTCTGACAGTCGCGGACACGGCCACCTATTTCTGTGCGAGAGCCGATACTGGTGCTGGTGGTTACCCGTTTAACTTGTGGGGCCCAGGCACCCTGGTCACCGTCTCCTCA | QQLEQSGGGLVQPEGSLTLTCTVSGFSWKYDYMCWVRQAPGKGLEWIVCIYGDSGSTWYASWVNGRFTISKTSSTTVTLQMASLTVADTATYFCARADTGAGGYPFNLWGPGTLVTVSS | GCCCTTGTGATGACCCAGACTCCAGCCTCCGTGTCTGAACCTGTGGGAGGCACAGTCATCATCAATTGCCAGGCCAGTCAGAGTATTGGTAGTAATTTAGCCTGGTATCAGCAGAAACCAGGGCAGCGTCCCAAGCTCCTGATGTATGCTGTTTCCAATCTGGCATCTGGGGTCCCATCGCGGTTCAGTGGCAGTGGATCTGGGACAGAGTTCACTCTCACCATCAGCGACCTGGAGTGGGCCGATGCTGCCACTTATTACTGTCAGGGCGGTGCGGTTTTTCATGATGTTGGTTTCGGCGGAGGGACCGAGGTGGTGGTCAAACGT | ALVMTQTPASVSEPVGGTVIINCQASQSIGSNLAWYQQKPGQRPKLLMYAVSNLASGVPSRFSGSGSGTEFTLTISDLEWADAATYYCQGGAVFHDVGFGGGTEVVVKR | This paper |
| M8a-7 IgG | Mouse | CAGGCTTACCTGCAGCAGTCTGGCGCTGAAATGGTTCGACCTGGCGCCTCCGTGAAGATGAGCTGTAAAGCCAGCGGCTACACCTTCAACAACTACAACATGCACTGGGTCAAGCAGACCCCTAGCCAGGGACTCGAGTGGATCGGCGGCTTTTACCCCGGCAATGACGATACCGCCTACAGCCAGAAGTTCAAGGGCAAAGCCACACTGACCGTGGACAAGAGCAGCAGCACCGCCTTTATGCACCTGAGCAGCCTGACCAGCGAGGACAGCGCCGTGTACTTTTGTGCCAGAAGCCTGGGCCGCTACTACGCCATGGATTATTGGGGCCAGGGCACCAGCGTGACCGTTTCT | QAYLQQSGAEMVRPGASVKMSCKASGYTFNNYNMHWVKQTPSQGLEWIGGFYPGNDDTAYSQKFKGKATLTVDKSSSTAFMHLSSLTSEDSAVYFCARSLGRYYAMDYWGQGTSVTVS | GACATCGTGCTGACACAGAGCCCTGCTTCTCTGGCCGTGTCTCTGGGACAGAGAGCCACCATCAGCTGTAGAGCCAGCGAGAGCGTGGACGACTTCGGCATCAGCTACATGAACTGGTTCCAGCAGAAGCCCGGCCAGACACCTAAGCTGCTGATCTACGGCGCCAGCAATCAAGGCAGCGGAGTGCCTGCCAGATTTTCTGGCTCTGGCAGCGGCACCGACTTCAGCCTGAACATTCACCCCATGGAAGAGGACGACCCCGCCATGTACTTTTGCCAGCAGAGCAAAGAGGTGCCCTACACCTTTGGCGGAGGCACCAAGCTGGAAATCAAG | DIVLTQSPASLAVSLGQRATISCRASESVDDFGISYMNWFQQKPGQTPKLLIYGASNQGSGVPARFSGSGSGTDFSLNIHPMEEDDPAMYFCQQSKEVPYTFGGGTKLEIK | Fan, et al., Immunity, 2022 |

**Table S2.** Cryo-EM data collection, refinement and validation statistics

| | M8b-A10 in complex with SARS-2 spike (EMDB 48347) (PDB 9ML4) | M8b-B8 in complex with SARS-2 spike (EMDB 48348) (PDB 9ML5) | M8b-C9 in complex with SARS-2 spike (EMDB 48349) (PDB 9ML6) | M8b-C10 in complex with SARS-2 spike (EMDB 48350) (PDB 9ML7) |
|---|---|---|---|---|
| **Data collection and processing** | | | | |
| Magnification | 105,000x | 45,000x | 105,000x | 105,000x |
| Voltage (kV) | 300 | 200 | 300 | 300 |
| Electron exposure (e–/Å$^2$) | 60 | 60 | 60 | 60 |
| Defocus range (μm) | -1 to -3 | -1 to -3 | -1 to -3 | -1 to -3 |
| Pixel size (Å) | 0.832 | 0.869 | 0.832 | 0.832 |
| Symmetry imposed | C1 | C1 | C1 | C1 |
| Initial particle images (no.) | 950,884 | 576,218 | 1,159,077 | 1,663,789 |
| Final particle images (no.) | 134,713 | 148,375 | 132,671 | 165,304 |
| Map resolution (Å) | 3.3 | 3.4 | 3.5 | 3.2 |
| FSC threshold | 0.143 | 0.143 | 0.143 | 0.143 |
| **Refinement** | | | | |
| Initial model used (PDB code) | 7UZD, 7SC1 | 7UZD, 7SC1 | 7UZD, 7SC1 | 7UZD, 7SC1 |
| Model composition | | | | |
| Non-hydrogen atoms | 29,480 | 27,822 | 29,751 | 25,846 |
| Protein residues | 3,717 | 3,491 | 3,750 | 3,259 |
| Ligands | 58 | 49 | 45 | 35 |
| *B* factors (Å$^2$) | | | | |
| Protein | 167.3 | 125.0 | 90.0 | 124.5 |
| Ligand | 145.1 | 120.0 | 97.2 | 128.3 |
| R.m.s. deviations | | | | |
| Bond lengths (Å) | 0.003 | 0.003 | 0.003 | 0.004 |
| Bond angles (°) | 0.603 | 0.575 | 0.610 | 0.630 |
| Validation | | | | |
| MolProbity score | 1.7 | 1.8 | 1.9 | 2.1 |
| Clashscore | 8.3 | 7.1 | 9.7 | 10.4 |
| Poor rotamers (%) | 1.2 | 1.4 | 1.7 | 2.5 |
| Ramachandran plot | | | | |
| Favored (%) | 96.7 | 96.2 | 96.9 | 96.3 |
| Allowed (%) | 3.3 | 3.8 | 3.1 | 3.7 |
| Disallowed (%) | 0 | 0 | 0 | 0 |

**Table S3.** X-ray data collection and refinement statistics

| | M8b-B1 Fab in complex with SARS-2 RBD (PDB 9ML8) | M8b-C9 Fab in complex with SARS-2 RBD (PDB 9ML9) |
|---|---|---|
| **Data collection** | | |
| Space group | C2 | C2 |
| Cell dimensions | | |
| *a*, *b*, *c* (Å) | 175.1, 173.4, 129.9 | 108.2, 127.2, 75.4 |
| α, β, γ (°) | 90, 116.6, 90 | 90, 127.7, 90 |
| Resolution (Å) | 2.40 - 39.14 (2.40 - 2.44) | 2.59 - 37.99 (2.59 - 2.71) |
| $R_{merge}$ | 0.172 (0.989) | 0.082 (0.983) |
| $I / \sigma I$ | 7.3 (2.3) | 11.7 (1.7) |
| Completeness (%) | 98.5 (99.1) | 98.5 (94.5) |
| Redundancy | 7.2 (7.4) | 7.2 (7.1) |
| **Refinement** | | |
| Resolution (Å) | 2.4 | 2.6 |
| No. reflections | 132,912 | 24,650 |
| $R_{work}$ / $R_{free}$ | 0.223/0.247 | 0.226/0.257 |
| No. atoms | 20,452 | 4,967 |
| Protein | 19,550 | 4,916 |
| Ligand/ion | 147 | 30 |
| Water | 755 | 23 |
| *B*-factors (Å$^2$) | 55.2 | 73.9 |
| Protein | 55.4 | 73.9 |
| Ligand/ion | 93.2 | 91.8 |
| Water | 43.6 | 61.0 |
| R.m.s. deviations | | |
| Bond lengths (Å) | 0.002 | 0.003 |
| Bond angles (°) | 0.54 | 0.55 |
| Ramachandran plot | | |
| Ramachandran favored (%) | 97.8 | 97.5 |
| Ramachandran allowed (%) | 2.2 | 2.5 |
| Ramachandran outliers (%) | 0 | 0 |
| Validation | | |
| Molprobity score | 1.2 | 1.4 |
| Clashscore | 3.9 | 5.8 |
| Rotamer outliers (%) | 0.4 | 0.6 |

*Values in parentheses are for highest-resolution shell.
